# Physical interactions between Gsx2 and Ascl1 regulate the balance between progenitor expansion and neurogenesis in the mouse lateral ganglionic eminence

**DOI:** 10.1101/794511

**Authors:** Kaushik Roychoudhury, Joseph Salomone, Shenyue Qin, Masato Nakafuku, Brian Gebelein, Kenneth Campbell

## Abstract

The Gsx2 homeodomain transcription factor is required to maintain neural progenitor identity in the lateral ganglionic eminence (LGE) within the developing ventral telencephalon, despite its role in upregulating the neurogenic factor Ascl1. How Gsx2 maintains cells as progenitors in the presence of a pro-differentiation factor is unclear. Here, we show that Gsx2 and Ascl1 are co-expressed in dividing subapical progenitors within the LGE ventricular zone (VZ). Moreover, we show that while Ascl1 misexpression promotes neurogenesis in dorsal telencephalic progenitors that do not express Gsx2, co-expression of Gsx2 with Ascl1 inhibits neurogenesis in these cells. To investigate the mechanisms underlying this inhibition, we used a cell-based luciferase assay to show that Gsx2 reduced the ability of Ascl1 to activate target gene expression in a dose-dependent and DNA binding-independent manner. Yeast 2-hybrid and co-immunoprecipitation assays revealed that Gsx2 physically interacts with the basic-Helix-Loop-Helix (bHLH) domain of Ascl1, and DNA binding assays demonstrated that this interaction interferes with the ability of Ascl1 to form homo- or heterodimers with E-proteins such as Tcf3 on DNA. To further assess for *in vivo* molecular interactions between these transcription factors within the telencephalon, we modified a proximity ligation assay for embryonic tissue sections and found that Ascl1:Gsx2 interactions are enriched within VZ progenitors, whereas Ascl1:Tcf3 interactions predominate in basal progenitors. Altogether, these findings suggest that physical interactions between Gsx2 and Ascl1 limit Ascl1:Ascl1 and Ascl1:Tcf3 interactions, and thereby inhibit Ascl1-dependennt neurogenesis and allow for progenitor expansion within the LGE.

## Introduction

Throughout evolution, the mammalian telencephalon, including both the cerebral cortex and the striatum (a.k.a. caudate-putamen), has enlarged dramatically, as compared to more caudal central nervous system (CNS) regions. This expansion has been made possible, at least in part, through a major increase in the number of telencephalic progenitors during embryogenesis (reviewed in Wilsch-Bräuninger et al., 2016). Along the rostral-caudal axis of the CNS, most neural progenitors in the ventricular zone (VZ) divide at the ventricular (i.e. apical) surface and have thus been termed apical progenitors (APs) (Wilsch-Bräuninger et al., 2016, Fish et al., 2008). APs comprise radial glia that undergo a process called interkinetic nuclear migration, wherein the soma migrates to the apical surface to undergo M-phase. The ventricular surface area thereby represents a limiting factor in this developmental process, and thus, a secondary progenitor population has arisen in the telencephalon at the basal extent of the VZ, termed basal progenitors (BPs) (Smart, 1976, Fish et al., 2008). The BPs comprise the subventricular zone (SVZ) and can expand through 1-2 rounds of cellular divisions before undergoing a terminal symmetrical neuronal division, thereby doubling neuronal output (Haubensak et al., 2004, Noctor et al., 2004). Recently, a new intermediate progenitor was identified in the VZ of the ganglionic eminences, including both the medial (MGE) and lateral ganglionic eminence (LGE). These dividing cells, which possess aspects of radial glia, have been termed subapical progenitors (SAPs) and give rise to the BPs (Pilz et al., 2013). Hence, SAPs are akin to the neural progenitors of the outer SVZ of the cerebral cortex of higher mammals and primates (Hansen et al., 2010, Fietz et al., 2010). Altogether, the APs and SAPs in the VZ, as well as the BPs in the SVZ allow for significant expansion of neuronal numbers in the mammalian telencephalon from rodents to primates. However, the molecular mechanisms that regulate their generation and differentiation remain unclear.

The LGE gives rise to striatal projection neurons, which comprise the vast majority of neurons in the striatum as well as olfactory bulb interneurons and a subset of amygdalar interneurons known as intercalated cells (ITCs) (Deacon et al., 1994, Olsson et al., 1995, Olsson et al., 1998, Wichterle et al., 2001, Waclaw et al., 2010). These neuronal subtypes arise from two defined progenitor domains within the LGE; the dorsal (d)LGE which produces the olfactory bulb interneurons and ITCs, and the ventral (v)LGE that generates the striatal projection neurons (Yun et al., 2001, Stenman et al., 2003, Waclaw et al., 2010). The homeodomain protein Gsx2 (a.k.a. Gsh2) is expressed by neural progenitors of the LGE (Szucsik et al., 1997, Toresson et al., 2000) with high levels of expression defining dLGE progenitors whereas moderate levels define those in the vLGE (Yun et al., 2001). Genetic lineage studies have shown that *Gsx2*-expressing LGE progenitors ultimately give rise to both neurons and glia (Kessaris et al., 2006, Fogarty et al., 2007, Qin et al., 2016). Accordingly, analysis of *Gsx2* mutants revealed an essential role for this transcription factor in generating each of the above-mentioned LGE-derived neuronal subtypes (Toresson et al., 2000, Corbin et al., 2000, Yun et al., 2001, Waclaw et al., 2009, Waclaw et al., 2010, Kuerbitz et al., 2018). Gsx2 is likely to regulate the generation of these neuronal subtypes through the upregulation of the proneural basic helix-loop-helix (bHLH) factor Ascl1 (Toresson et al., 2000, Corbin et al., 2000, Yun et al., 2001, Waclaw et al., 2009, Wang et al., 2009). In fact, some Gsx2^+^ LGE progenitors co-express Ascl1 (Yun et al., 2003, Wang et al., 2013). However, previous studies have indicated that maintained expression of Gsx2 limits the ability of LGE (Pei et al., 2011, Méndez-Gómez and Vicario-Abejón, 2012) and postnatal SVZ cells (López-Juárez et al., 2013) to differentiate, Furthermore, *Gsx2* mutant mice, which exhibit reduced levels of Ascl1, have LGE progenitors that precociously generate oligodendrocyte precursor cells (Chapman et al., 2013). Thus, Gsx2 primes LGE progenitors for neurogenesis by upregulating Ascl1, but these cells remain as undifferentiated progenitors that undergo further expansion until Gsx2 expression is down-regulated (Pei et al., 2011). In this study, we describe how molecular interactions between Gsx2 and Ascl1 within subsets of LGE progenitors may impact the choice between progenitor expansion versus neurogenesis.

## Results

### Gsx2 and Ascl1 define the progression of LGE progenitor maturation

Previous studies revealed Gsx2 is upstream of the neurogenic factor Ascl1 in LGE VZ progenitors (Toresson et al., 2000, Corbin et al., 2000, Yun et al., 2001, Waclaw et al., 2009, Wang et al., 2009), thus specifying a neurogenic potential within this population. Moreover, a portion of Gsx2-expressing LGE progenitors co-express Ascl1 (Yun et al., 2003, Wang et al., 2013), suggesting progenitors progress from Gsx2-only VZ cells to Ascl1-only VZ/SVZ cells via a transitional state in which both factors are co-expressed. To better define this progenitor progression, we examined Gsx2 and Ascl1 expression within LGE progenitors between E11.5 and E15.5, which represents the early neurogenic period when secondary progenitors (e.g. BPs) are established (Bhide, 1996, Pilz et al., 2013). Gsx2/Ascl1 double-labeled cells occurred in a “salt and pepper” fashion throughout the LGE VZ but were generally absent in the most apical cells (Fig. 1A-C and Suppl Fig. 1). At early stages of neurogenesis, Ascl1^+^ cells comprise only about 1/3 of Gsx2^+^ LGE VZ cells, however, the vast majority of Asc1^+^ cells co-express Gsx2 (Fig. 1A-C, H, I and Suppl. Fig. 1A). The proportion of double-labeled cells in the vLGE, which contributes to striatal neurogenesis extensively between E11.5 and E15.5 (Waclaw et al., 2009, Kelly et al., 2018), was dynamic. Indeed, the number of Gsx2^+^Ascl1^+^ double-labeled cells as a percentage of Gsx2^+^ cells peaked at E13.5 (Fig. 1H and Suppl. Fig. 1A-C), which correlates with the establishment of the proliferative SVZ (Bhide, 1996). Moreover, by E15.5, the percentage of double-labeled vLGE cells as a proportion of Ascl1^+^ cells, fell to roughly half of that seen at E11.5 (Fig. 1I), resulting in more Ascl1-only (i.e. SVZ BPs) cells during peak striatal neurogenesis. In contrast, the proportion of double-labeled cells as a ratio of either Gsx2 or Ascl1 did not change much over time in the dLGE (Fig. 1H, I and Suppl. Fig. 1A-C), where robust neurogenesis occurs after E15.5 and into postnatal stages (Hinds, 1968, Tucker et al., 2006, Waclaw et al., 2006, López-Juárez et al., 2013).

**Figure 1.**
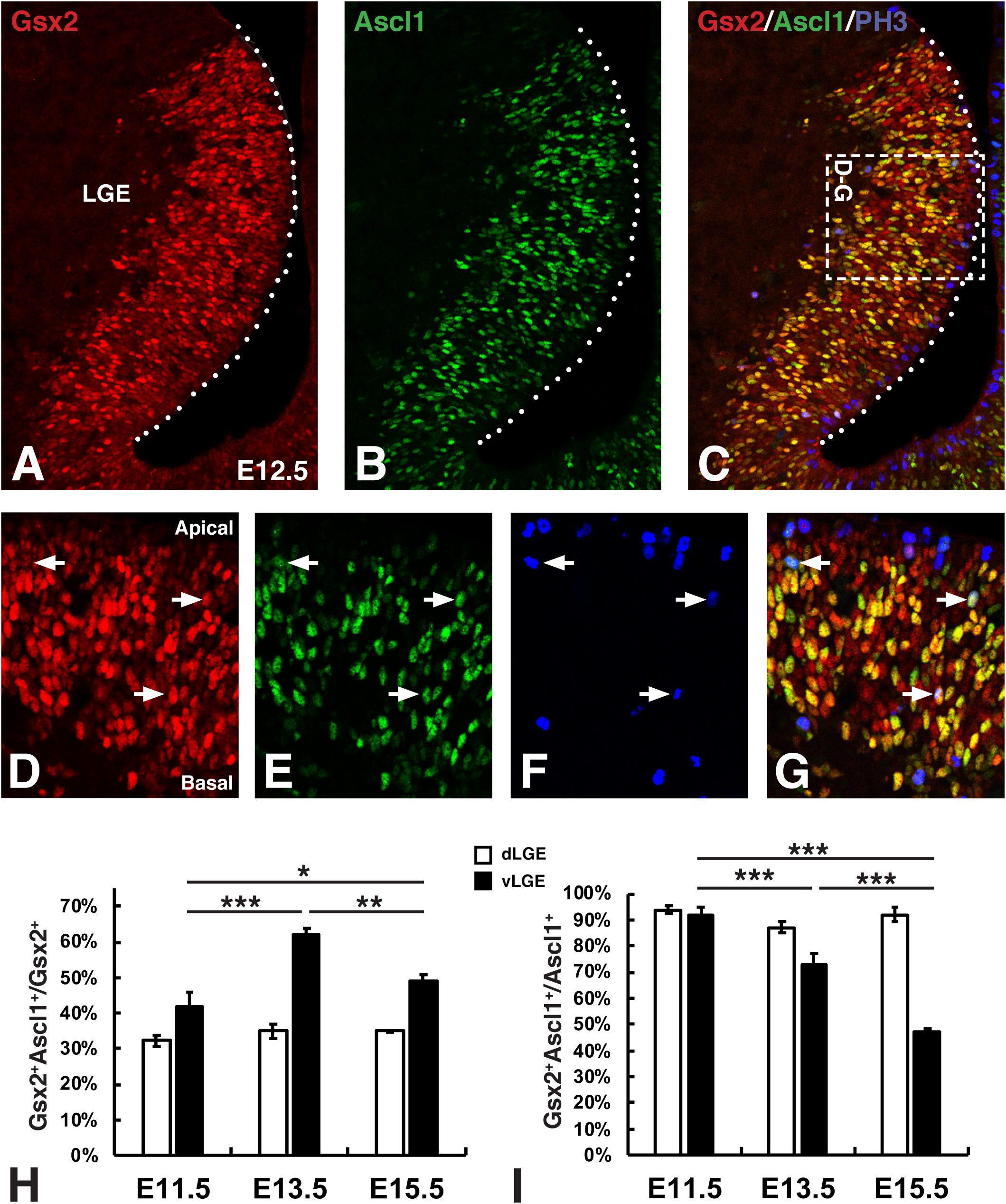
Gsx2 and Ascl1 co-expression marks LGE subapical progenitors (SAPs). Triple immunohistochemistry for Gsx2 (**A, D**), Ascl1 (**B, E**) and phosphohistone 3 (PH3) (**C, F, G**) in the E12.5 LGE. Box in **C** has been rotated 90 counterclockwise for the images in **D**-**G**. Note that most APs (i.e. PH3^+^ cells at the apical surface indicated by dotted lines in **A**-**C** or positioned at the top of **D**-**G**) express low or undetectable levels of either Gsx2 or Ascl1 (**C, G**). In contrast, PH3^+^ cells at abventricular positions (i.e. SAPs) within the VZ frequently co-localize Gsx2 and Ascl1 (**D-G**). Quantification of Gsx2^+^Ascl1^+^ co-expressing cells in the dLGE (open bars) versus vLGE (black bars) as either a ratio of the Gsx2^+^ (**H**) or Ascl1^+^ (**I**) cells from the embryonic stages shown in Suppl. Fig. 1. Data shown in **H** and **I** represent the mean ± SD (n=3). Note, a small but significant (p<0.05) difference was detected in the Gsx2^+^Ascl1^+^/Ascl1^+^ cells of the dLGE at E13.5 (**H**). One-way ANOVA was performed between the dLGE or the vLGE data at each embryonic stage with a Tukey posthoc. *p<0.05, **p<0.01 and ***p<0.001.

To examine how the dynamic expression of Gsx2 and Ascl1 correlates with LGE progenitor subtypes (i.e. APs, SAPs and BPs), we triple labeled with phosphohistone 3 (PH3) to identify dividing progenitors in M phase. Most PH3-positive M phase cells lined the ventricular (i.e. apical) surface, and thus represent APs (Fig. 1C). These apical dividing cells typically expressed no, or low levels of Gsx2 and in rare instances, they expressed low levels of Ascl1 (Fig. 1D-G). In contrast, the vast majority of subapical PH3^+^ LGE cells were found to co-express both Ascl1 and Gsx2 (arrows in Fig. 1D-G). The Ascl1^+^ cells that divide at abventricular positions within the VZ were previously described as a unique progenitor population within the VZ, termed subapical progenitors (SAPs), which are dependent on Ascl1 for their appearance in the LGE VZ as well as for the normal generation of proliferative progenitors in the SVZ (i.e. BPs) (Castro et al., 2011, Pilz et al., 2013). Thus, Gsx2 and Ascl1 co-expression appears to be a defining feature of LGE SAPs and together these factors may regulate the expansion potential of these intermediate progenitors.

### Gsx2 expression overrides Ascl1-induced neurogenesis in the mouse telencephalon

To better understand the impact of Gsx2 and Ascl1 co-expression in telencephalic progenitors, we used a mouse *Foxg1^tTA^* transgenic system to either misexpress each factor individually or together in the dorsal telencephalon (Hanashima et al., 2002, Waclaw et al., 2009, Ueki et al., 2015). We previously used this system to show that Gsx2 misexpression ventralizes dorsal telencephalic progenitors and induces an LGE fate, including the upregulation of Ascl1 (Waclaw et al., 2009, Chapman et al., 2013). Moreover, Gsx2 misexpression maintained telencephalic cells in a neural progenitor state with a concomitant reduction in neurogenesis (Pei et al., 2011).

To determine the impact of Ascl1 misexpression on dorsal telencephalic progenitors, we analyzed E12.5 *Foxg1^tTA^; tet-O-Ascl1* and *Foxg1^tTA^* control embryos for Gsx2 and Ascl1 as well as neuronal differentiation markers. As expected, *Foxg1^tTA^; tet-O-Ascl1* embryos expressed Ascl1 throughout the dorsal-ventral extent of the telencephalon with no change in Gsx2 expression, which stops at the pallio-subpallial boundary (arrowheads in Fig. 2A and B). A previous study by Fode et al. (2000) showed that ectopic expression of Ascl1 in dorsal telencephalic progenitors drives ventral (i.e. LGE) identity similar to Gsx2 misexpression (Waclaw et al., 2009) along with the specification of GABAergic neuronal phenotypes. Using our misexpression system, we found that ectopic Ascl1 induced a significant increase in neurogenesis in the dorsal telencephalon as marked by ß-III-tubulin (Tubb3) and doublecortin (Dcx) (Fig. 2F, I, J), as compared to controls (Fig. 2E, I, J). Because the dense staining for both of these immature neuron markers made cellular analysis difficult, we measured the thickness of the pallial region expressing Tubb3 and Dcx (short bars) as a ratio of total pallial thickness (longer bars) in Fig. 2E-H. Consistent with our previous publication (Pei et al., 2011), we found that Gsx2 misexpression, despite upregulating Ascl1 (Fig. 2D), resulted in reduced Tubb3 and Dcx expression in the dorsal telencephalon (Fig. 2 H-J). Not only was the ratio of the Tubb3/Dcx-positive staining domain to total pallial thickness significantly reduced, the amount of staining for either neuronal marker was greatly diminished as compared to both Ascl1 misexpressing embryos and control embryos (compare Fig. 2H with 2E and 2F). Since the levels of Gsx2 are likely to be significantly higher than the amount of up-regulated Ascl1 in *Foxg1^tTA^; tet-O-Gsx2* embryos, the imbalance could favor neural progenitor maintenance over neurogenesis. To test this idea, we generated *Foxg1^tTA^; tet-O-Ascl1; tet-O-Gsx2* embryos to simultaneously express both factors in dorsal telencephalic progenitors (Fig. 2C). As anticipated, Gsx2 and Ascl1 levels are similarly increased, but like in Gsx2 misexpressing embryos, neurogenesis was reduced in embryos misexpressing both Gsx2 and Ascl1 (Fig. 2G-J). Thus, co-expression of Gsx2 and Ascl1 in dorsal progenitors severely limits Ascl1-driven neurogenesis, consistent with Gsx2/Ascl1 co-expression in LGE SAPs allowing for progenitor expansion at the expense of neurogenesis.

**Figure 2.**
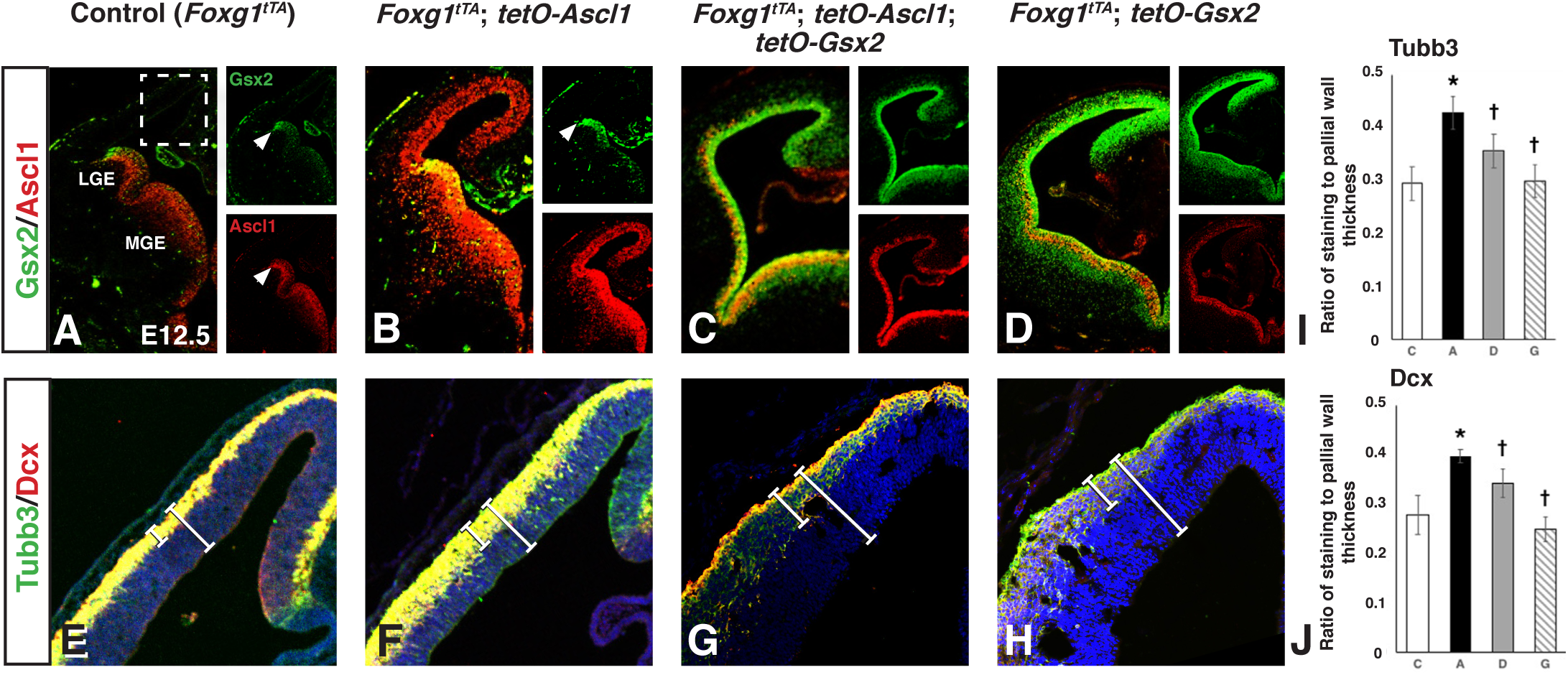
Gsx2 inhibits Ascl1-driven neurogenesis in a transgenic misexpression assay. Coronal sections through the telencephalon of E12.5 control (i.e. *Foxg1^tTA^*) (**A, E**), Ascl1 misexpressing (i.e. *Foxg1^tTA^; tetO-Ascl1*) (**B, F**), Gsx2 and Ascl1 misexpressing (i.e. *Foxg1^tTA^; tetO-Ascl1; tetO-Gsx2*) (**C, G**) and Gsx2 misexpressing (i.e. *Foxg1^tTA^; tetO-Gsx2*) (**D, H**) embryos. **B**) Misexpression of Ascl1 throughout the telencephalon did not alter Gsx2 expression, which remained in the subpalium, stopping at the pallio-subpallial boundary (indicated by arrowheads) as in controls (**A**). Misexpression of Ascl1 within the dorsal telencephalon did, however, lead to an increase in Tubb3 and Dcx staining (compare **F** with **E**). Quantification was made by measuring the Tubb3/Dcx-positive cortical staining (smaller white bar) and represented as a ratio of the total pallial wall (larger white bar). Data presented in **I** (Tubb3) and **J** (Dcx) represent mean ± SD for each genotype (C, control, n= 3; A, Ascl1 misexpression, n= 3; D, Gsx2 and Ascl1 misexpression, n= 3; G, Gsx2 misexpression, n= 3). Misexpression of Gsx2 alone upregulated Ascl1 throughout the telencephalon (**D**) and reduced neurogenesis (**H)**, which was similar to misexpression of both Gsx2 and Ascl1 (**C, G**). One-way ANOVA was performed between the data from C, A, D and G embryos with a Tukey posthoc. *p<0.05 as compared to control (C) and †p<0.01 as compared to Ascl1 (A) misexpressing embryos.

### Gsx2 inhibits Ascl1-mediated gene expression in a dose-dependent manner

Ascl1, like other bHLH transcription factors, activates target gene expression by binding as a dimer to palindromic DNA sequences referred to as E-boxes (consensus CANNTG sequence) (Johnson et al., 1992, Henke et al., 2009). Homodimeric or heterodimeric complexes can bind these targets, but for class II, tissue specific bHLHs like Ascl1, heterodimerization with more widely expressed class I bHLHs, termed E-proteins, is often important for target activation (Massari and Murre, 2000, Henke et al., 2009). To examine whether Gsx2 interferes with Ascl1-mediated gene expression, we developed an *in vitro* luciferase assay in *Drosophila* S2 cells. We utilized *Drosophila* S2 cells to eliminate the confounding effects of Gsx2 upregulation of endogenous Ascl1 in mammalian cell lines. In this assay, we used a portion of the *Drosophila achaete* promoter (Jafar-Nejad et al., 2003), which contains 3 distinct E-box binding sites fused to a luciferase cassette, termed *Ac-Luc* (Fig. 3A). This construct shows low basal activity and interestingly, when Ascl1 alone is added, no significant increase in luciferase was detected (Fig. 3B). The addition of Daughterless (i.e. *Drosophila* E-protein) alone resulted in a modest increase (approx. 5-fold) in luciferase, whereas co-expression of Ascl1 and E-protein resulted in a roughly 33-fold increase in luciferase expression (Fig. 3B). Titrating in increasing levels of Gsx2 resulted in a stepwise reduction of luciferase expression (Fig. 3B), supporting the notion that Gsx2 limits the transcriptional activity of Ascl1. In contrast, Gsx2 failed to alter the ability of Daughterless (i.e. E-protein homodimers) to activate luciferase in this cell-based assay (Fig. 3C). Given that Gsx2 is known to have repressor functions (Winterbottom et al., 2010, 2011), it is possible that Gsx2 directly binds DNA to repress the *achaete* promoter. To test this idea, we mutated the asparagine (N) at amino acid position 253 to an alanine (A) in Gsx2’s homeodomain (Gsx2^N253A^) which abrogates DNA binding (Fig. 3D-F). Importantly, we found that titrating in increasing levels of Gsx2^N253A^ also resulted in a significant reduction in luciferase activity, with the exception of the lowest concentration tested (Fig. 3B). These findings suggest that the ability of Gsx2 to reduce Ascl1-mediated gene expression is largely independent of Gsx2 DNA binding.

**Figure 3.**
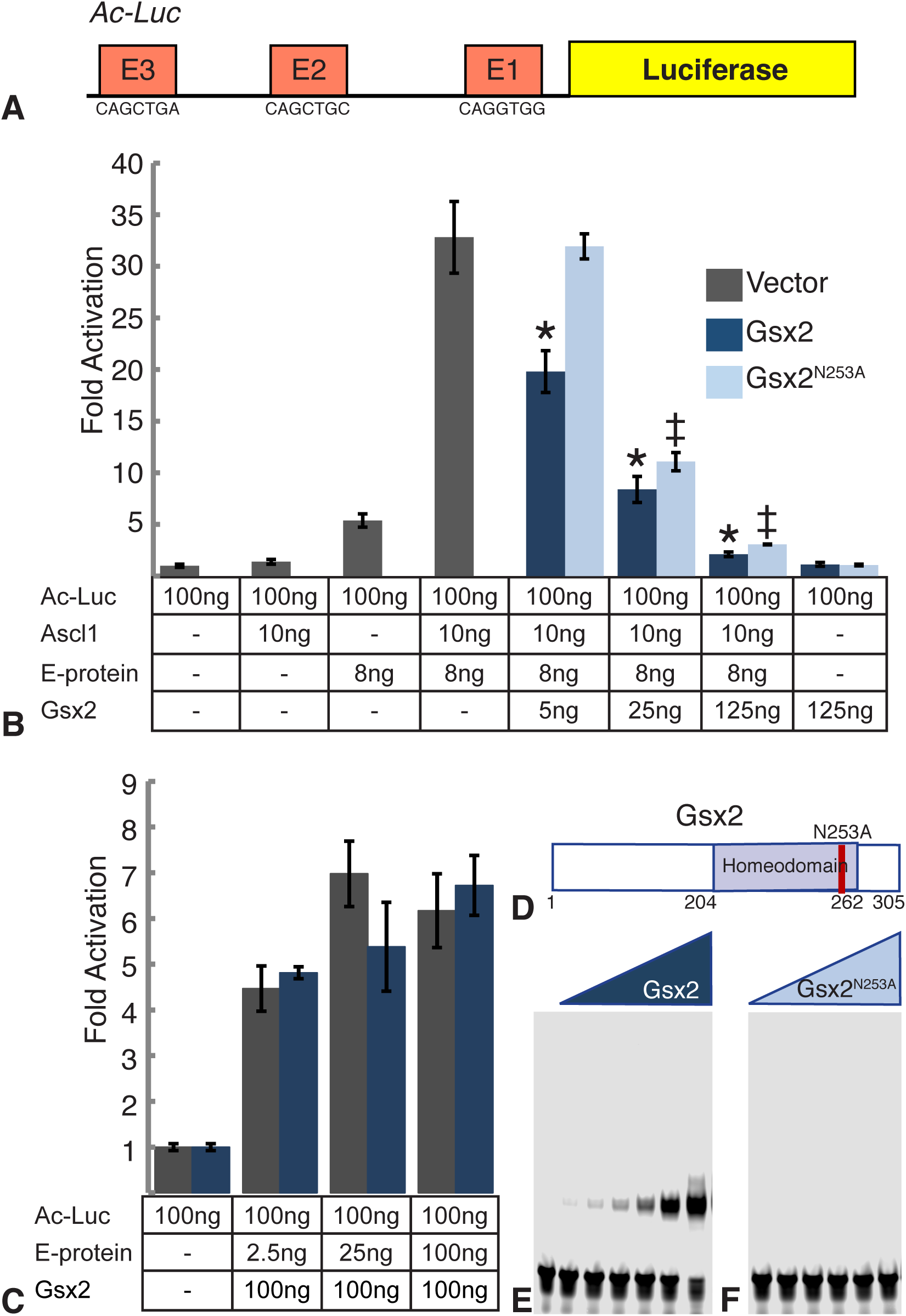
Gsx2 interferes with Ascl1-mediated reporter activation independent of its ability to bind DNA. **A**) Schematic of the Luciferase reporter construct used in **B** and **C**. 420 bp from the promoter of the *Drosophila acheate* gene contains 3 E-box sequences that can be bound by Ascl1 (Supplemental Figure 2). **B**) 100ng of the *Ac-Luc* reporter was co-transfected with 10ng Ascl1, 8ng of the *Drosophila* E-protein (Daughterless), and the indicated amount of Gsx2. Values represent fold activation over the *Ac-Luc* reporter added alone. Effects of co-transfecting an empty pAC5.1 expression vector, Gsx2 wild type, and Gsx2 DNA binding mutant (N253A) are shown. **C**) 100ng of *Ac-Luc* reporter was co-transfected with the indicated amount of *Drosophila* E-protein with 100ng of wild type Gsx2. Values represent fold activation over the *Ac-Luc* reporter added alone. All conditions were performed in triplicate, data represent means ± SD. In **B**, a one-way ANOVA was performed between vector, Gsx2 and Gsx2^N253A^ was performed with a Tukey posthoc. * indicates significant difference (p<0.01) between transfection of empty vector and Gsx2 WT. ‡ indicates significant difference(p<0.01) between transfection of empty vector and Gsx2^N253A^. In **C**, a student’s t-test was performed between vector and Gsx2. **D**) Schematic of the Gsx2 protein indicating the homeodomain, and the position of the amino acid mutated to disrupt DNA binding. Equimolar amounts of Gsx2 (**E**) and Gsx2^N253A^(**F**) were added to probes containing a predicted high affinity Gsx2 binding site. Note, the complete loss of DNA binding with the Gsx2^N253A^ protein.

### Gsx2 interacts with Ascl1 in LGE progenitors

The mechanism by which Gsx2 limits Ascl1’s activity is unclear. One possibility is that Gsx2 might bind Ascl1 at the protein level and interfere with its function. To test this hypothesis, we performed a yeast 2-hybrid interaction assay, using either Ascl1 or Olig2, another bHLH factor expressed in subsets of LGE progenitors (Takebayashi et al., 2000, Chapman et al., 2013) as *prey* and Gsx2 as *bait*. In this system, we found that Gsx2 robustly interacts with Ascl1 but not Olig2 (Fig 4A, B). Moreover, consistent with the results from the luciferase assay showing that Gsx2 did not alter E-protein homodimer-mediated gene activation (Fig. 3C), we did not observe interactions between Gsx2 and Tcf3 (i.e. mouse E-protein) in the yeast 2-hybrid assay (Fig. 4C). Interestingly, the interaction with Ascl1 appears to be rather specific to Gsx2, as yeast 2-hybrid experiments using Ascl1 as *prey* and Gsx1, a close homolog of Gsx2, as *bait* failed to show an interaction (Fig. 4D). To test whether Gsx2 indeed interacts with Ascl1 in the mouse forebrain, we performed co-immunoprecipitation (IP) assays on lysates from E12.5 telencephalons using antibodies specific for either Gsx2 or Ascl1. Subsequent Western blot analysis revealed that Ascl1 co-IPs with Gsx2 and vice versa (Fig. 4E), suggesting that Gsx2 and Ascl1 physically interact within LGE progenitors of the mouse embryonic telencephalon.

**Figure 4.**
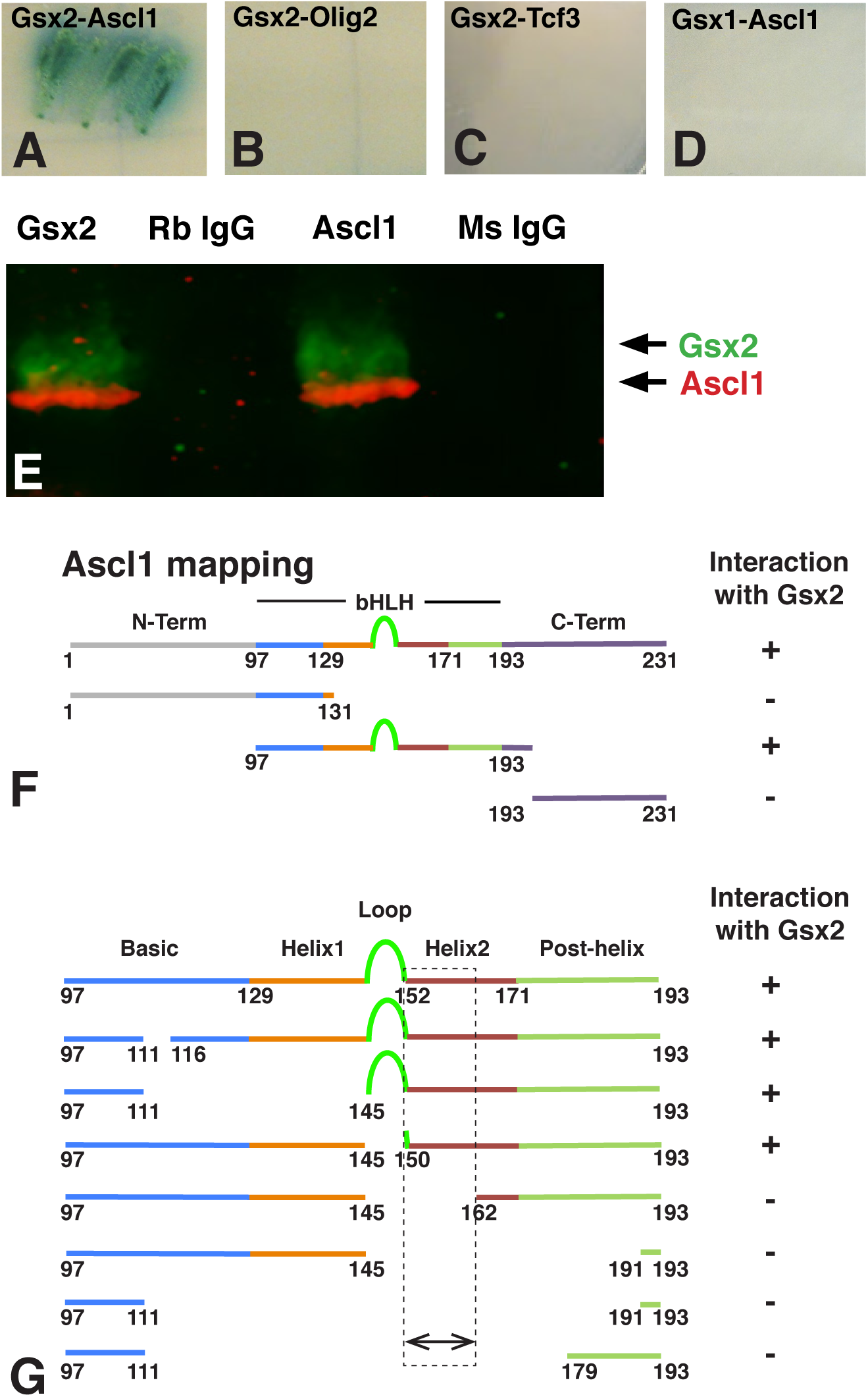
Gsx2 physically interacts with the bHLH domain of Ascl1 in the mouse telencephalon. Yeast 2-hybrid experiments using either Gsx2 as bait and Ascl1 as prey (**A**), Gsx2 as bait and Olig2 as prey (**B**), Gsx2 as bait and Tcf3 as prey (**C**) or Gsx1 as bait and Ascl1 as prey (**D**). Note, robust reporter gene expression (i.e. *β*-gal) was only observed in the Gsx2-Ascl1 experiment (**A)**. **E**) Co-IP experiments using lysates from E12.5 mouse telencephalon. Pulling down Gsx2 with a rabbit antibody and blotting with a mouse Ascl1 antibody showed association of Ascl1 with Gsx2. Conversely, pulling down Ascl1 and blotting with a Gsx2 antibody showed association of these two proteins in the embryonic mouse telencephalon. Rabbit IgG and mouse IgG were used as controls for the Gsx2 and Ascl1 pull downs, respectively. **F**) Using truncated portions of Ascl1 (e.g. N-Term, bHLH and C-Term) in a yeast 2-hybrid assay, we found that only the bHLH domain of Ascl1 interacts with Gsx2. **G**) Further deletion mapping studies using the yeast 2-hybrid assay showed that the second helix (amino acids 150-162) of Ascl1 is required for interactions with Gsx2.

### Gsx2 binds the second helix of the Ascl1 bHLH domain

To determine which domain within Ascl1 interacts with Gsx2, we generated and tested deletion constructs in the yeast 2-hybrid assay. Given that Gsx2 interferes with the neurogenic function of Ascl1, we hypothesized that Gsx2 may interact with a portion of Ascl1 necessary for DNA binding and/or forming transcription complexes. To test this possibility, we performed yeast 2-hybrid assays with constructs that either express the Ascl1 N-terminus (Ascl1^1-131^), bHLH domain (Ascl1^97-193^), or C terminus (Ascl1^193-231^) and found that only the bHLH domain mediated interactions with Gsx2 (Fig. 4F). To further map the sub-domain of the Ascl1 bHLH that interacts with Gsx2, we used site-directed mutagenesis to create a series of small deletions (Fig. 4G). Of these domains, the basic region and first helix of the bHLH are necessary for DNA binding, while the second helix is involved in protein dimerization with other bHLH proteins including itself (homodimers) as well as with E-proteins such as Tcf3 (heterodimers) (Johnson et al., 1992, Massari and Murre, 2000, Nakada et al., 2004, Henke et al., 2009). Interestingly, only mutations inside or spanning the second helix of Ascl1’s bHLH domain disrupted Gsx2 binding (Fig. 4G). This result suggests that Gsx2 may compete with Ascl1 and/or E-proteins (e.g. Tcf3) in the formation of homo- and heterodimers.

### Gsx2:Ascl1 interactions inhibit Ascl1 homo- and heterodimer complexes from binding DNA target E-boxes

Given that Gsx2 physically interacts with the same portion of the Ascl1 bHLH domain that mediates dimer formation with other bHLH proteins, we hypothesized that Gsx2 interferes with Ascl1’s ability to form homo- or heterodimers with E-proteins on its target DNA, the E-box. To test this idea, we used purified Gsx2, Ascl1, and Tcf3 (i.e. E-protein) in electromobility shift assays (EMSAs). To select an appropriate DNA probe, we tested the 3 identified E-box sequences from the *Ac-Luc* construct for Ascl1:Ascl1 homodimer, Ascl1:Tcf3 heterodimer, and Tcf3:Tcf3 homodimer binding (Suppl. Fig. 2). From these studies, we found that the E2-box sequence (Fig. 3A), which most closely matches the known consensus Ascl1 binding motif (Castro et al., 2011), mediated the most robust formation of both Ascl1:Ascl1 and Ascl1:Tcf3 complexes (Suppl. Fig. 2A-I). To examine the effect of Gsx2 on homo- and heterodimer formation on the E2 site, we performed EMSAs with increasing concentrations of Gsx2. Note, all experiments were performed with the Gsx2^N253A^ protein, which is unable to bind DNA, but capable of inhibiting Ascl1-induced gene expression (Fig. 3B and 3F). Titrating in increasing levels of Gsx2^N253A^ efficiently disrupted Ascl1 homodimer formation on the E2 sequence (Fig. 5B, C). As it is technically difficult to determine the relative concentrations of Ascl1 or Tcf3 within LGE progenitors, we performed EMSAs with ratios of Ascl1 and Tcf3 ranging from Ascl1:Tcf3 (E-protein) = 32:1, 1:2 and 1:12 (Fig. 5D, F, H). In EMSAs performed with a high ratio of Ascl1 relative to Tcf3 (32:1), only Ascl1:Ascl1 homodimers and Ascl1:Tcf3 heterodimers were detected and increasing the amount of Gsx2^N253A^ preferentially diminished Ascl1 homodimer formation (Fig. 5D, E). At a 1:2 Ascl1:Tcf3 ratio, we predominantly detect Ascl1:Tcf3 heterodimers with only weak Tcf3:Tcf3 homodimers observed (Fig 5F, lane 16). Under these conditions, adding increasing amounts of Gsx2^N253A^ did not significantly alter Ascl1:Tcf3 heterodimer formation (Fig. 5F, G). However, a slight increase in Tcf3 homodimer binding was observed (Fig. 5F, G), likely as a result of Gsx2 binding Ascl1, and thereby freeing increasing amounts of Tcf3 that can form homodimer complexes on DNA. EMSAs performed with a 1:12 Ascl1 to Tcf3 ratio and increasing amounts of Gsx2^N253A^ showed no effect on Tcf3 homodimer formation but a significant reduction of Ascl11:Tcf3 heterodimers (Fig. 5H, J). Finally, no effect was observed by increasing levels of Gsx2^N253A^ on the formation of Tcf3 homodimers in the absence of Ascl1 (Fig. 5J, K), consistent with Gsx2 neither physically interacting with Tcf3 (Fig. 4C) nor inhibiting Tcf3 homodimer-induced luciferase activation (Fig. 3C). Altogether, these data support a model in which Gsx2 physically interacts with the Ascl1 bHLH domain, and thereby interferes with the formation of Ascl1 homodimers and Ascl1-Tcf3 heterodimers required to bind DNA. Hence, the inhibition of Ascl1-mediated gene expression by Gsx2 (Fig. 3B) is likely to result from its ability to interfere with Ascl1 binding to E-boxes in the *Ac-Luc* construct.

**Figure 5.**
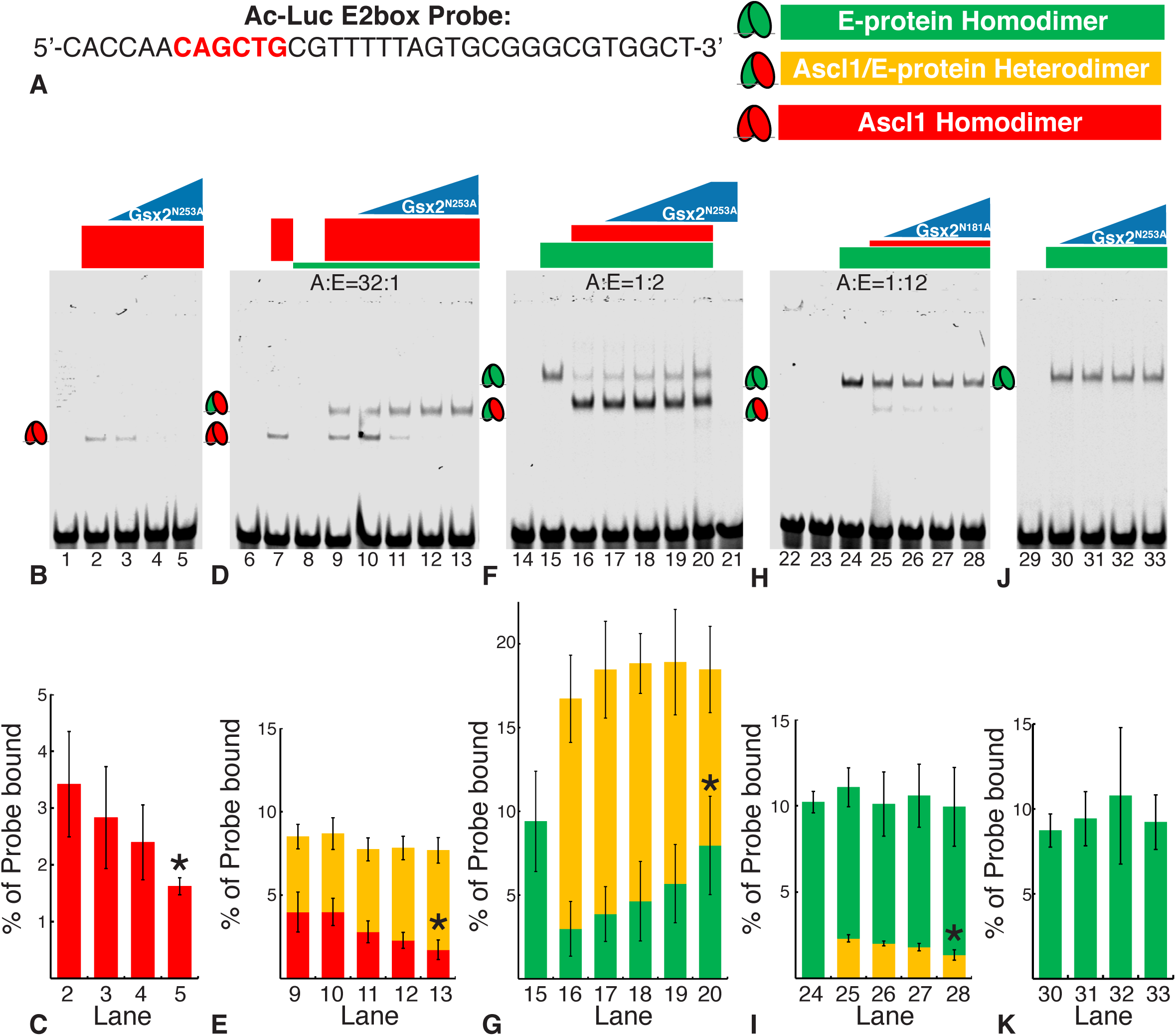
Gsx2 interferes with Ascl1 homodimer and heterodimer binding to an E-box DNA sequence. **A**) Sequence of the E-box (highlighted in red) probe used in lanes 1-33 of EMSAs shown in **D**-**K**. **B**) Gsx2 DNA binding mutant, Gsx2^N253A^, is titrated in increasing amounts from 0 to 80 pmoles in samples containing a constant 2.5 pmoles of Ascl1. **C**) Percentage of probe bound by Ascl1-Ascl1 homodimers (red bars) in lanes 2-5. **D**) Ascl1 (2.5pmoles) and E-protein (0.08pmoles) were added in a 32 to 1 ratio in each lane (9-13) with increasing levels of Gsx2^N253A^ from 0 to 80 pmoles. **E**) Percentage of probe bound by Ascl1-Ascl1 homodimers (red) and Ascl1-E47 heterodimers (yellow) in lanes 9-13. **F**) Ascl1 (0.15 pmoles) and E-protein (0.3 pmoles) were added in a 1 to 2 ratio in each lane (16-20) with increasing levels of Gsx2^N253A^ from 0 to 80 pmoles. **G**) Percentage of probe bound by Ascl1-E47 heterodimers (yellow) and E47-E47 homodimers (green) in lanes 15-20. **H**) Ascl1 (0.026 pmoles) and E-protein (0.31 pmoles) were added in a 1 to 12 ratio in each lane (25-28) with increasing levels of Gsx2^N253A^ from 0 to 80 pmoles. **I**) Percentage of probe bound by Ascl1-E47 heterodimers (yellow), and E47-E47 homodimers (green) in lanes 24-28. **J**) E-protein (0.3 pmoles) was added to each lane (30-33) with increasing levels of Gsx2^N253A^ from 0 to 80 pmoles. **K**) Percentage of probe bound by E47-E47 homodimers (green) in lanes 30-33. Each EMSA was performed in triplicate, and data in **C** through **K** represent mean ± SD with the intensity of bands representing each complex normalized to total probe intensity. A student’s t-test was performed between the no Gsx2^N253A^ condition and the maximum Gsx2^N253A^ condition, *p<0.05.

### Gsx2 competes with the E-protein, Tcf3 to bind to Ascl1

Ascl1 can homodimerize as well as heterodimerize with Tcf3 isoforms (e.g. E-proteins E12/E47) via protein-protein interactions (Johnson et al., 1992, Nakada et al., 2004, Henke et al., 2009). Since bHLH homo- and heterodimer formation require the second helix and this helix also interacts with Gsx2, we assessed whether Gsx2 competes with Tcf3 to interact with Ascl1. Note, we did not observe interactions between Tcf3 and Gsx2 in a yeast 2-hybrid assay (Fig. 4C). To investigate if Gsx2 competes with E-proteins to interact with Ascl1, we used a yeast 3-hybrid system in which the production of a third interfering protein is turned on or off using a methionine (met) inducible switch (Tirode et al., 1997). We made use of the “met off” system to test the impact of either Tcf3 on Gsx2:Ascl1 interactions or Gsx2 on Ascl1:Tcf3 interactions. In the absence of Tcf3 (+ met), Ascl1 (*prey*) interacted with Gsx2 (*bait*) (Fig 6A). In the presence of Tcf3 (- met), however, the Ascl1:Gsx2 interaction was largely abrogated (Fig 6B). Thus, a typical Ascl1 interacting partner, Tcf3, can interfere with Gsx2:Ascl1 interactions. Interestingly, the converse experiment using Tcf3 as *bait* and Ascl1 as *prey* in the presence or absence of Gsx2 showed that Gsx2 is also capable of abrogating Ascl1:Tcf3 interactions (Fig. 6C, D). Hence, these data are consistent with Ascl1 being able to interact with either Gsx2 or Tcf3 but not both at the same time.

**Figure 6.**
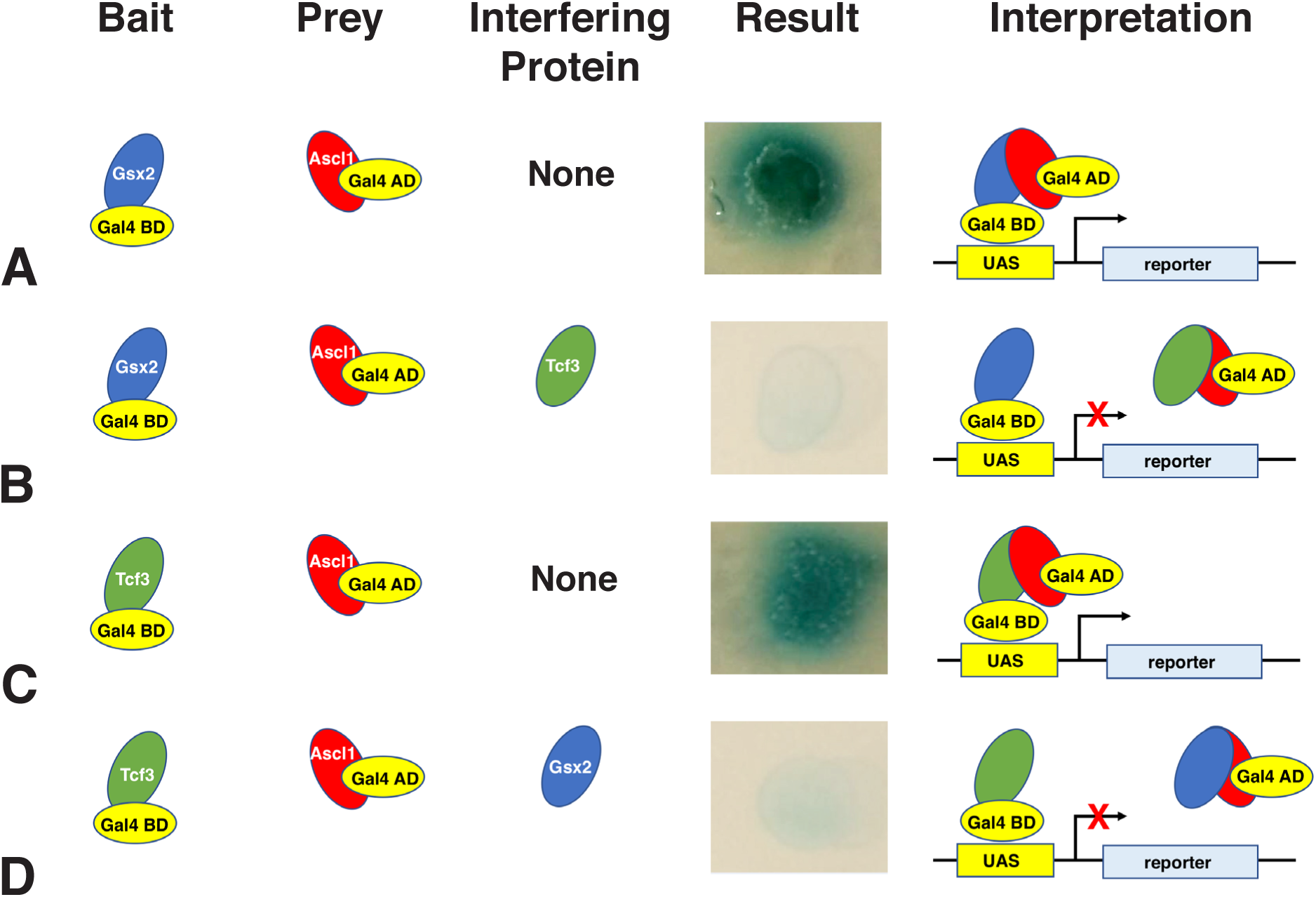
Gsx2 and Tcf3 compete for molecular interactions with Ascl1 in a yeast 3-hybrid assay. The yeast 3-hybrid assay utilizes an interfering protein which is capable of interacting with the bait or the prey to disrupt their interaction. **A**) Using Gsx2 as bait and Ascl1 as prey with no interfering protein results in reporter (*β*-gal) expression, while addition of Tcf3 (E-protein) as an interfering protein disrupts the interaction as shown in **B**. **C**) Likewise, with Tcf3 as bait and Ascl1 as prey and no interfering protein, *β*-gal expression is activated, which is disrupted by addition of Gsx2 as the interfering protein as seen in **D**.

### Spatial localization of Gsx2:Ascl1 and Ascl1:Tcf3 interactions within the LGE

To further characterize Gsx2:Ascl1 protein interactions *in vivo*, we adapted a <u>p</u>roximity ligation <u>a</u>ssay (PLA) protocol from cultured cells (Söderberg et al., 2006, Bagchi et al., 2015) for use with paraformaldehyde fixed mouse embryonic forebrain tissue sections. PLA uses two secondary antibodies conjugated to short oligonucleotides, to which mutually complementary oligonucleotides are ligated *in situ*. If two proteins are adjacent to or bound to each other and the antibodies are within 40nm (Bagchi et al., 2015), the oligonucleotide tags prime a fluorescent-based polymerization reaction that results in repetitive loops using a rolling circle amplification model. We performed PLA to detect Gsx2:Ascl1 interactions in E12.5 telencephalons and found that there is widespread signal throughout the LGE VZ (Fig. 7A), which appears as single fluorescent dots at high power (Fig. 7B). Indeed, the spatial pattern of the PLA signal for Gsx2:Ascl1 interactions fits well with the double immunofluorescent staining for Gsx2 and Ascl1 (Fig. 1C). To demonstrate PLA signal specificity, we performed PLA for Gsx2 and Ascl1 in *Gsx2* knockout mice and found no detectable signal (Fig. 7C). As a further control, we utilized sections from embryos that either misexpress Gsx2 (*Foxg1^tTA^, tetO-Gsx2*) or Ascl1 *(Foxg1^tTA^, tet-O-Ascl1*). In embryos misexpressing Gsx2, we observed PLA signal in VZ progenitors throughout the dorsal-ventral aspect of the telencephalon (Fig. 7D). This finding is in accordance with the fact that Gsx2 upregulates Ascl1 in dorsal telencephalic progenitors (Waclaw et al., 2009) (see also Fig. 2D). In contrast, *Foxg1^tTA^; tet-O-Ascl1* embryos only showed a PLA signal in the ventral telencephalon (Fig. 7E), consistent with Gsx2 not being upregulated in the pallium of these embryos (Fig. 2B). Finally, we performed PLA for Gsx2 and Ascl1 in sections from the *Foxg1^tTA^; tet-O-Ascl1; tet-O-Gsx2* embryos and observed robust PLA signal within VZ progenitors throughout the dorsal-ventral aspect of the telencephalon (Fig 7F). Taken together, our data demonstrate that Gsx2 and Ascl1 physically interact within the LGE VZ progenitors that co-express these two factors.

**Figure 7.**
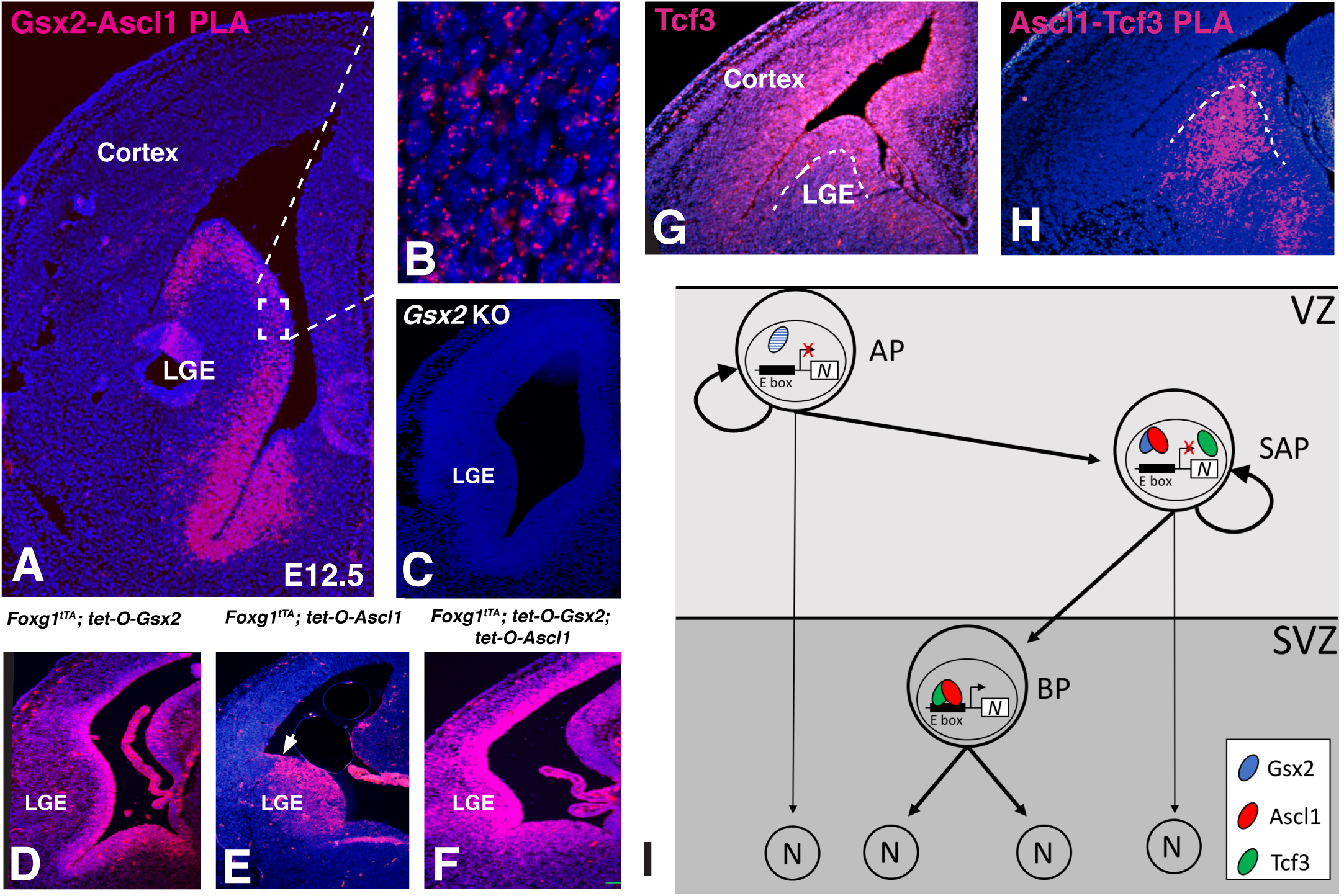
Proximity ligation assay (PLA) shows Ascl1:Gsx2 interactions and Ascl1:Tcf3 interactions in distinct portion of the LGE germinal zone. **A**) When PLA was performed using rabbit anti-Gsx2 and guinea pig anti-Ascl1 antibodies, strong signal (magenta) was detected in the LGE and septal VZ. **B**) High power magnification of the VZ region shows punctate signal associated with the DAPI-stained nuclei. **C**) The PLA signal between Gsx2 and Ascl1 antibodies was specific as no signal was detected in *Gsx2* knock-out (KO) tissue sections. **D**) In Gsx2 misexpressing embryos, PLA signal is expanded throughout the telencephalon as is the case for Gsx2 and Ascl1 expression. **E**) PLA signal is not expanded throughout the telencephalon in the Ascl1 misexpressing embryos (pallio-subpallial boundary indicated by arrow in **E**) as Gsx2 is not upregulated outside of the ventral telencephalon (see Fig. 2B). However, the PLA signal is intensified in the ventral telencephalon. **F**) Misexpression of both Gsx2 and Ascl1 leads to an increased PLA signal throughout the telencephalon. **G**) Immunostaining for Tcf3 protein in the E12.5 telencephalon shows staining throughout the germinal zones including both the VZ and SVZ. **H**) PLA using the goat anti-Tcf3 and guinea pig anti-Ascl1 antibodies shows signal in both the LGE VZ as well as the SVZ, with stronger signal in the latter region. The boundary between the VZ and SVZ is indicated by the dashed line in **G** and **H**. **I**) Schematic model showing LGE progenitor subtypes, with SAPs co-expressing Gsx2 and Ascl1 thus limiting Ascl1’s neurogenic function and allowing for progenitor expansion. Gsx2 expression is lost in BPs thus allowing Ascl1:Tcf3 heterodimers to direct neurogenesis. Note, Gsx2 was observed in some APs (indicated by dashed lines) but not together with Ascl1. In this model both APs and SAPs could undergo direct neurogenesis if Gsx2 was down-regulated (indicated by thin arrows).

Our results from the yeast 3-hybrid assay raise the possibility that competition between Gsx2 and Tcf3 to interact with Ascl1 may occur in LGE cells. Indeed, immunostaining for Tcf3 in E12.5 telencephalon showed broad staining, with the highest signal confined to the germinal regions including the VZ, where Gsx2 and Ascl1 are co-expressed (Fig 7G). Thus, there is extensive overlap between these transcription factors in the LGE, particularly in the VZ. To investigate Ascl1:Tcf3 interactions in the LGE, we performed PLA using Tcf3 and Ascl1 antibodies. Interestingly, strong PLA signal was predominantly detected within the SVZ of the LGE, while a weaker signal was observed in the VZ (Fig. 7H). Since Gsx2 is largely confined to the VZ, the reduced PLA signal for Ascl1:Tcf3 interactions suggests that Gsx2 competes with Tcf3 for Ascl1 interactions in the VZ. Taken together, our results suggest that Gsx2:Ascl1 interactions predominate in LGE VZ cells (i.e. SAPs) while neurogenic Ascl1:Tcf3 interactions characterize SVZ cells (i.e. BPs) (Fig. 7I).

## Discussion

In this study, we show that a subset of LGE VZ progenitors, namely the SAPs, co-express Gsx2 and Ascl1. Moreover, we demonstrate that Gsx2 limits Ascl1-driven neurogenesis, at least in part, through an interaction with its bHLH domain, thereby limiting Ascl1’s ability to form homo- and heterodimers, bind target DNA sequences, and activate gene expression. Finally, using PLA, we found that Gsx2:Ascl1 interactions occur predominantly in LGE VZ cells (presumably SAPs), while Ascl1:Tcf3 interactions largely occur in the BPs of the SVZ. Thus, interactions between Gsx2 and Ascl1 are likely to contribute to the expansion of LGE progenitors, particularly the SAPs, by delaying neurogenesis until Gsx2 is downregulated in the LGE SVZ BPs (see schematic in Fig. 7I). In this way, Gsx2 primes the neuronal potential of LGE progenitors by up-regulating Ascl1 expression, but also allows for sufficient progenitor (e.g. SAP) expansion to generate the proper number of neurons.

The co-expression of Gsx2 and Ascl1 within the LGE SAPs is in line with a number of previous findings in *Gsx2* and *Ascl1* mutants. Firstly, *Gsx2* null mutants lack a proliferative SVZ (i.e. BPs) in the LGE at early stages of neurogenesis, which recovers partially after the *Gsx1* family member is upregulated in the mutant (Toresson and Campbell, 2001). A similar loss of BPs was also recently reported in the septal region where *Gsx2* was conditionally inactivated (Qin et al., 2017), supporting the notion that Gsx2 is required to generate secondary progenitors (e.g. SAPs and BPs) in the ventral telencephalon. Interestingly, Ascl1 not only marks the LGE SAPs, but is required for their generation as well as the production of BPs (Pilz et al., 2013). Thus, given that Gsx2 is necessary for the normal expression of Ascl1 in LGE progenitors (Toresson et al., 2000, Corbin et al., 2000, Yun et al., 2001), it seems likely that in addition to the observed loss of BPs in the *Gsx2* mutant SVZ (Toresson and Campbell, 2001), SAPs would also be missing in these mutants. Taken together, co-expression of Gsx2 and Ascl1 both mark the LGE SAPs and regulate their development into BPs which subsequently contribute to LGE-derived neurons (see Fig. 7I).

The vLGE gives rise to the striatal projection neurons (Yun et al., 2001, Stenman et al., 2003) which are organized into two neurochemically and anatomically distinct compartments, termed the patch (a.k.a. striosome) and matrix (Graybiel and Ragsdale, 1978, Gerfen, 1992). The patch, which comprises only 15% of the striatum, is generated at early stages in the rodent (e.g. E10-12 in the mouse), whereas the matrix compartment occupies approximately 85% of the striatum and is populated at later stages (e.g. E13 and onward) (van der Kooy and Fishell, 1987, Johnston et al., 1990). This difference in timing and the larger size contribution of matrix projection neurons to the striatum correlates well with the appearance of the Ascl1^+^ SAPs in the mouse vLGE. For example, SAPs appear in the mouse forebrain between E12-14 and represent nearly half of all dividing LGE VZ progenitors by E16 (Pilz et al., 2013). Moreover, consistent with the finding that the *Ascl1* lineage undergoes an abrupt transition around E13, wherein *Ascl1*-expressing progenitors exhibit an expanded capacity for proliferation, as predicted if they progress through the SAP to BP expansion route (Kelly et al., 2018). Thus, this change in the *Ascl1* lineage correlates with the generation of the patch versus matrix compartments of the striatum and suggests that to generate the large numbers of matrix neurons, expansion through SAPs and BPs is likely necessary.

The co-expression of Gsx2 and the neurogenic factor Ascl1 in LGE SAPs presents a challenge for these cells to maintain progenitor status in the presence of a pro-differentiation factor. We previously showed that Gsx2 plays a role in maintaining progenitor identity within the LGE lineage (Pei et al., 2011). Hence, despite upregulating Ascl1, Gsx2 expressing telencephalic progenitors do not undergo rapid neuronal differentiation. In this study, we confirmed this finding, as well as compared and contrasted the ability of Ascl1 to induce neurogenesis in cells that do not express Gsx2 (i.e. the dorsal telencephalon) versus those that co-express both Ascl1 and Gsx2 (i.e. double mis-expressing embryos). Our findings reveal that even when expressing high levels of both Gsx2 and Ascl1, dorsal telencephalic progenitors show a pronounced decrease in neurogenesis to a level similar to those misexpressing Gsx2 only. Like Ascl1, the *Dlx* genes require Gsx2 for their correct expression in LGE progenitors (Toresson et al., 2000, Corbin et al., 2000, Yun et al., 2001, Wang et al., 2013). Both Ascl1 and Dlx factors have been shown to promote the differentiation of GABAergic neuronal phenotypes typical of the LGE (Yun et al., 2002, Long et al., 2009a, 2009b, Pla et al., 2018, Lindtner et al., 2019). Furthermore, loss of Gsx2 leads to an upregulation of the oligodendrocyte precursor cell marker Pdgfr*α* (Corbin et al., 2003) and concomitant precocious oligodendrocyte differentiation (Chapman et al., 2013, 2018). Thus, it appears that Gsx2 specifies a neurogenic potential in LGE progenitors by virtue of upregulating Ascl1 (and Dlx factors) but maintains the co-expressing cells as progenitors capable of expanding as SAPs and subsequently generating BPs.

Gsx1 is known to partially compensate for loss of *Gsx2* in LGE progenitors (Toresson and Campbell, 2001, Yun et al., 2003). However, previous misexpression studies showed that Gsx1 promotes neuronal differentiation in the dorsal telencephalon, whereas Gsx2 (see e.g. Fig. 2) inhibits neurogenesis (Pei et al., 2011). Similar to Gsx2, Gsx1 is capable of upregulating Ascl1 when misexpressed in telencephalic progenitors (Pei et al., 2011, Chapman et al., 2013, 2018). Moreover, Gsx1 is required for the restoration of Ascl1 expression in LGE progenitors of *Gsx2* mutants (Toresson and Campbell, 2001, Yun et al., 2003). Our results, however, show that, at least in the yeast 2-hybrid assay, Gsx1 does not appear to interact with Ascl1. Thus, the observation that misexpression of Gsx1 leads to increased neurogenesis (Pei et al., 2011) is consistent with Gsx1 inducing Ascl1 expression, which in turn promotes neurogenesis.

Our data suggests that the manner in which Gsx2 limits Ascl1-driven neurogenesis in LGE progenitors is by disrupting its ability to form functional homo- and heterodimers and thereby decreasing DNA binding to E-box sequences. Formation of Ascl1:Ascl1 and Ascl1:E-protein dimers occurs though interactions between amino acids in the second helix of the bHLH and is independent of DNA binding (Massari and Murre, 2000, Nakada et al., 2004). Remarkably, Gsx2 specifically interacts with this portion of the Ascl1 protein, and thus in cells that express Gsx2, Ascl1, and E-protein, such as the LGE SAPs, Gsx2 may limit the neurogenic capacity of Ascl1. Indeed, our 3-hybrid experiments support this notion. It is important to mention, however, that in addition to inhibiting Ascl1:Ascl1 and Ascl1:E-protein dimer binding target DNA, it is also possible that Gsx2:Ascl1 complexes may bind novel DNA target sequences, which could also promote progenitor maintenance. Unlike the case for Ascl1, Gsx2 binding its target DNA sequence was not disrupted by titrating in higher amounts of Ascl1 (Suppl. Fig. 3) suggesting that the region of Gsx2 that interacts with Ascl1 is outside of the homeodomain. So far, we do not know the specific residues of Gsx2 that interact with Ascl1, nor do we know whether Ascl1 binding affects other potential binding partners of Gsx2 or if it leads to unique target gene selection. Further studies will be needed to address these questions.

Although Ascl1 is a well-known neurogenic factor, it has also been implicated in progenitor maintenance. In fact, Imayoshi et al. (2013) showed that *Ascl1* gene and protein expression levels oscillate at moderate levels in neural progenitors, and only when Ascl1 expression levels increase and become sustained does it drive neuronal differentiation. Thus, it may be that the interaction with Gsx2 limits the ability of the moderate oscillatory Ascl1 levels from regulating genes involved in neuronal differentiation, and it is only when Gsx2 expression is downregulated (e.g. in BPs) that Ascl1 expression stabilizes and reaches a high enough level to regulate gene expression required for neurogenesis. Alternatively, Ascl1 may utilize a Gsx2-independent mechanism to reach a high and sustained level of expression in LGE progenitors which at a certain threshold could overcome the limiting interactions with Gsx2. To date, there are no reports of *Gsx2* gene or protein oscillations in LGE progenitors, however, it is not clear that the issue has been addressed. It also should be noted that Castro et al. (2011) identified two distinct sets of gene targets for Ascl1; one group that contributes to neurogenesis and another that regulates progenitor proliferation. Our EMSA data (Fig. 5) using the “E2” E-box sequence suggests that Ascl1:Ascl1 homodimers may be more sensitive to Gsx2 disruption than Ascl1:Tcf3 heterodimers. Although it is unclear whether the homo-versus heterodimer complexes have distinct preferred DNA binding sites, and by extension, regulate genes differentially, it could be that Gsx2 co-expression favors Ascl1:Tcf3 over Ascl1:Ascl1 complexes in progenitor cells aiding in the maintenance of progenitor identity.

Finally, we have modified the PLA technique (Söderberg et al., 2006, Bagchi et al., 2015) to examine the LGE cells in which Gsx2:Ascl1 and/or Ascl1:E-protein interactions occur in tissue sections. This technology allows for the identification of cells in which two proteins are within 40 nm of each other and thus, are likely in direct physical contact (Bagchi et al., 2015). This represents one of the only reports we are aware of using the PLA technique to visualize protein:protein interactions in brain sections. The great advantage of this technology is that we can test for regional localization of different partner proteins within complex tissues. Indeed, using this technique, we were able to detect Gsx2:Ascl1 interactions in LGE VZ cells, whereas Ascl1:E-protein interactions predominate in the LGE SVZ. These findings support the progenitor lineage model proposed in Figure 7I, wherein Gsx2:Ascl1 interactions in LGE SAPs limit Ascl1’s neurogenic potential and thereby permit further expansion of neuronally specified progenitors either as SAPs or as BPs. In contrast, because BPs lack Gsx2, Ascl1:E-protein or Ascl1:Ascl1 interactions could promote symmetric neurogenic divisions to generate neurons. Thus, the differential protein-protein interactions between Gsx2:Ascl1 and Ascl1:Tcf3 within tissues provide a novel mechanism in which the choice of transcription factor partner ultimately dictates distinct cellular responses: Gsx2:Ascl1 interactions allow for continued progenitor expansion, whereas Ascl1:Tcf3 interactions promote cell cycle exit and subsequent neurogenesis.

## Materials and methods

### Animals

All experiments using mice were approved by the Institutional Animal Care and Use Committee (IACUC) of the Cincinnati Children’s Hospital Research Foundation and were conducted in accordance with US National Institutes of Health guidelines. The previously described transgenic mice used included *Foxg1^tTA^* (Hanashima et al., 2002), *tet-O-Gsx2* (Waclaw et al., 2009), *tet-O-Ascl1* (Ueki et al., 2015) and *Gsx2^RA/+^* (Waclaw et al., 2009). Maintenance and genotyping of animals and embryos was performed as described (Waclaw et al., 2009, Ueki et al., 2015). For misexpression studies, *Foxg1^tTA^* males were crossed with either *tet-O-Gsx2*, *tet-O-Ascl1* or *tet-O-Gsx2; tet-O-Ascl1* transgenic females. Additionally, *Gsx2^RA/+^* mice were intercrossed to generate *Gsx2^RA/RA^* null embryos. Day of vaginal plug detection was deemed E0.5.

### Immunohistochemistry

Embryos were fixed for 3 hrs (E11.5) or 6 hrs (E12.5) or overnight (E15.5) at 4°C in 4% PFA before washing three times in PBS and cryopreservation in 30% sucrose. Embryos were sectioned coronally at 12 µm thickness on a cryostat and slides were stored at −20°C until used.

Fluorescent immunohistochemistry on embryonic brain sections was performed as previously described (Waclaw et al., 2009). Primary antibodies were used at the following concentrations: guinea pig anti-Ascl1, 1:10,000 (provided by Jane Johnson, UT Southwestern, Dallas TX); mouse anti-Ascl1, 1:500 (BD Pharmingen); Guinea Pig anti-Dcx, 1:3000 (Millipore); rabbit anti-Gsx2, 1:3000 (Toresson et al., 2000); mouse anti-phosphohistone 3 (PH3), 1:500 (Cell Signaling); goat anti-Tcf3, 1:500 (Abcam) and rabbit anti-Tubb3, 1:1000 (Covance). Secondary antibodies used were: donkey anti-rabbit conjugated to Alexa 594 or Alexa 647 (Jackson ImmunoResearch), 1:200; donkey anti-guinea pig conjugated to Alexa 488 or Alexa 647 (Jackson ImmunoResearch), 1:200; donkey anti-goat IgG conjugated to Alexa 594 (Jackson ImmunoResearch), 1:200; goat anti-mouse IgG1 conjugated to Alexa 568 (Invitrogen), 1:500. Because all of the *tet-O* transgenes have an IRES-EGFP (Waclaw et al., 2009, Ueki et al., 2015) no Alexa 488 conjugated secondary antibodies were used on these sections. Slides were counterstained with DAPI for 10 minutes, coverslip-mounted with Fluoromount G and dried overnight at room temperature. In certain cases, slides were coverslipped with Fluormount G containing DAPI. Stained slides were imaged using confocal microscopy using a Nikon A1 LSM system with a GsAsP solid state laser.

### Quantification of immunostainings

For quantification of Gsx2 and Ascl1 cellular co-expression in the embryonic LGE, a box spanning the Gsx2-expressing VZ was drawn in the dLGE and vLGE, respectively, and Gsx2^+^ single-, Ascl1^+^ single- and Gsx2^+^Ascl1^+^ double-labeled cells were manually counted. 4 LGEs per embryo and 3 embryos at each timepoint were quantified. Statistical significance was determined by one-way ANOVA with the Tukey HSD post hoc test.

To quantify the effects of misexpression of Ascl1 alone, Gsx2 alone or Ascl1 and Gsx2 together on the staining of Dcx and Tubb3 in the dorsal telencephalon we measured the thickness of the immunostaining (indicated by the representative smaller white bar in Fig. 2) at ten locations spanning the dorsal pallium for each hemisphere as a ratio of the total mean pallial wall thickness at the same ten locations (indicated by the representative larger white bar in Fig. 2). Three embryos for each genotype were analyzed. Statistical significance was determined by one-way ANOVA with the Tukey HSD post hoc test.

### Plasmids, molecular cloning, and site-directed mutagenesis

Details describing all plasmids generated for this study can be found in Suppl. Table 1, and all primer sequences used for PCR-based cloning can be found in Suppl. Table 2. The following plasmids were purchased and used in this study: the pGBKT7, pGADT7 and pBRIDGE yeast vectors (Clontech); the pET14b bacterial protein expression vector (Novagen); and the pAc5.1 *Drosophila* expression vector (Invitrogen). The *Ac-*Luciferase and Da-pAc5.1 used for luciferase assays (Jafar-Nejad et al., 2003), and the E47 (isoform of Tcf3) amino acids 430-648 in pET14b used for bacterial protein expression (Zhang et al., 2019) were previously described. PCR reactions were performed using either Clonamp Hi-fidelity polymerase mix (Clontech) or Accuzyme DNA polymerase. Ligation reactions were performed using T4 DNA ligase or In-Fusion HD (Clontech). Site-directed mutagenesis was performed using inverse PCR followed by In-Fusion ligation. All DNA clones and mutations were verified by DNA sequencing.

### Yeast 2- and 3-hybrid assays

Yeast 2-hybrid assays were performed using the Matchmaker Gold yeast 2-hybrid system (Clontech 630489), and one of two previously described methods with minor modifications (<u>Clontech Yeast Protocols Handbook</u>). For the Gsx2:Ascl1, Gsx1:Ascl1, Gsx2:Tcf3 and Gsx2:Olig2 interaction assays, a ‘Mate and Plate’ (Letteboer et al., 2008) approach was taken. Briefly, a Y2H-Gold strain of *Saccharomyces cerveciae (Sc)* was freshly made competent by the lithium acetate method (Clontech Yeast Protocols Handbook) and the *bait* construct (Gsx2 or Gsx1 in the pGBKT7 vector) was transformed by PEG/DMSO and heat shock, as described (Clontech Yeast Protocols Handbook). The Y187 strain of *Sc* was similarly made competent and transformed with *prey* constructs (Ascl1, Olig2 or Tcf3 in the pGADT7 vector). Positive colonies were obtained using single-dropout nutritional auxotropy on SD-Trp and SD-Leu plates respectively. *Bait* and *prey* colonies were mixed and co-cultured overnight in 2x YPDA, mating confirmed by microscopy, and 2-hybrids were selected on double dropout (DDO) plates (SD-Leu,-Trp). Bait:prey interactions were determined on DDO + Aerobasidin A + X-*α*-Gal plates. Presence of blue colonies indicated positive interaction. The interactions were confirmed by patching the colonies in SD-Leu,-Trp, -His, -Ade, +ABA, +X-*α−*Gal plates.

To study interactions between full-length Gsx2 and Ascl1 deletion mutants, a ‘co-transformation’ approach was taken, as described (Chien et al., 1991). Briefly, both *bait* and *prey* constructs were co-transformed into the Y187 strain and two hybrids were selected on DDO plates. The rest of the assay was performed as described above.

Yeast 3-hybrid assays were performed essentially as described (Tirode et al., 1997). Briefly, a fresh competent Y187 strain of *Sc* was co-transformed with Ascl1 in the pGADT7 vector and either Gsx2-MCS1; E47 MCS2, E47 MCS1; Gsx2 MCS2, Gsx2 MCS1 or Tcf3 MCS1 constructs, as described above. Three hybrids were selected on DDO plates, bait-prey interaction determined on DDO/X/A plates and interference determined on SD-Trp, -Leu, -Met, XaGal, ABA, or the same plates + Met. Interference was confirmed by patching colonies on SD-Trp, -Leu, -His, -Ade, XaGal, ABA with or without Met.

### Immunoprecipitation and Western blots

Immunoprecipitations (IPs) were performed as described (Lin and Lai et al., 2017) with minor variations. Briefly, telencephalons were dissected from E12.5 mouse embryos in ice cold PBS. Each telencephalon was homogenized on ice with a plastic pestle in a microcentrifuge tube in 400 µl of modified RIPA buffer (50mM Tris pH7.4, 150mM NaCl, 1% Tx-100) supplemented with protease inhibitor cocktail (PIC) (Sigma P8340). Lysates were centrifuged at 15,000xG for 10 minutes at 4°C, and the supernatant was collected in a fresh, prechilled tube. 2 µl of rabbit anti-Gsx2 antibody or 5 µl of mouse anti-Ascl1 antibody, was added and the tube was rotated end over end at 4°C overnight. A 10 µl slurry of Protein G agarose (Sigma P7700) in ice cold RIPA buffer was prepared, added to the lysate and incubated a further two hours. The beads were separated from the liquid by centrifugation at 5000xG for five minutes at 4°C and washed five times in ice cold RIPA buffer with PIC. The IP’d proteins were eluted in Glycine-HCl buffer (pH 3) and denatured for 10 minutes at 70°C in SDS loading buffer containing *β−*mercaptoethanol.

For Western Blot analysis, samples were separated on a 4-20% acrylamide gel using a Tris-Glycine buffer. Gels were transferred to Nitrocellulose membranes with Tris/Glycine/Methanol transfer buffer in a semi-dry blotter (Atto Corp, Tokyo, Japan). Membranes were washed 3 times in TBS and blocked with 0.5% Casein (Sigma C7087) for 1 hour at room temperature. The membranes were then cut and probed with primary antibodies (rabbit anti-Gsx2 or mouse anti-Ascl1) in TBS + 0.1% Tween20 (TBST) overnight at 4°C with gentle rocking, washed thrice in TBST, and subsequently incubated with 1:15,000 LiCor secondary antibodies at room temperature for 2 hours. The membranes were then washed, dried and imaged in a LiCor Fc fluorescence imager at 700 and 800nm.

### Luciferase reporter assays

Luciferase assays were carried out in *Drosophila* S2 cells. S2 cells were cultured in HyClone media (Fisher Scientific). 6×10^5^ cells were cultured in 12-well plates for 24 hours prior to transfection. Each well was transfected with a total of 0.3 μg of DNA (100ng *Ac-luciferase* reporter, 2 ng *copia Renilla* and the indicated amount of expression constructs) using Effectene Transfection Reagent (Qiagen). Empty pAc5.1 was added to bring the total DNA to 0.3 μg per well. Cells were harvested 36 hours after transfection, lysates were isolated and luciferase activity was measured via Promega Dual Luciferase Assay kit. All firefly luciferase values were normalized to Renilla Luciferase values to normalize for variations in transfection efficiency, and each experiment was performed in triplicate. Results are reported as fold expression over the Ac-Luc reporter alone.

### Protein Purification and DNA probe design

His-tagged proteins were purified from BL21 cells via Nickel chromatography as described previously (Uhl et al., 2010). Fragments of the following *Mus musculus* proteins were used: Ascl1 amino acids 92-231 containing the bHLH domain, E47 (isoform of Tcf3 gene) amino acids 430-648 containing the bHLH domain, Gsx2 amino acids 167-305 containing the homeodomain, and Gsx2^N253A^ amino acids 167-305 containing the mutated homeodomain. Protein concentrations were determined via Bradford assay and confirmed via SDS-PAGE and Gel Code Blue stain (Thermo-fisher). Probes were generated as described previously by annealing a 5’-IRDye® 700 labeled oligo (IDT) to the oligos listed (below) and filling in with Klenow (Uhl et al., 2016).

### Electrophoretic Mobility Shift Assays (EMSAs)

EMSAs were performed as previously described using native PAGE (Uhl et al., 2010, 2016). For EMSAs in Supplemental Figure 2, 0.04 pmoles of the indicated probe was added to 20µL binding reactions. For EMSAs in Figure 5 and Supplemental Figure 3, 0.68 pmoles of probe was added to each 20µL binding reaction. To facilitate exchange between bHLH dimerization partners in experiments where Ascl1 and E47 were used, all binding reactions were incubated at 37°C for 30 minutes before addition of the probe. For binding reactions testing the ability of Gsx2 ^N253A^ to alter Ascl1:Ascl1 or Ascl1:E47 binding, the Ascl1 and Gsx2^N253A^ proteins were mixed first, incubated at 37°C for 30 minutes, and then E47 was added and samples were incubated at 37°C for an additional 30 minutes. Probes were added and samples incubated at room temperature for 10 minutes. For EMSAs shown in Figure 5, the number of pmoles of protein added in each lane is indicated in Supplementary Table 3. Quantification of EMSAs was performed using LI-COR Biosciences’s Image Studio software by calculating the total probe in each lane and the percentage of probe that is bound by either homodimer or heterodimer complexes. The following oligos were used for the EMSAs:

*Ac-Luc E1*: GTCACG**CAGGTG**GGATCCTAGTGCGGGCGTGGCT
*Ac-Luc E2*: CACCAA**CAGCTG**CGTTTTTAGTGCGGGCGTGGCT
*Ac-Luc E3*: GACAGG**CAGCTG**AAAATGTAGTGCGGGCGTGGCT
*Gsx2_site*: CAGAGTG**TAATTAA**CATTCAGTAGTGCGGGCGTGGCT

### Proximity ligation assay (PLA)

Duolink^®^ PLA kit components (Millipore Sigma, DUO92105) were used for performing PLA with a modified protocol. Positive (+ve) and negative (−ve) Rabbit and Goat PLA conjugated antibodies were purchased from Sigma. For the guinea-pig PLA antibody, the −ve PLA DNA tag was covalently attached to 100 µl of 1mg/ml of unconjugated donkey anti-guinea pig IgG Fab (Jackson Immuno Research) using the Duolink^®^ in situ probemaker minus kit (Millipore Sigma, DU092010) as per manufacturer’s instructions. PFA fixed cryostat sections (four per slide, one of each genotype) were dried immediately after cryosectioning and re-hydrated in 1xPBS for five minutes. The tissue sections were then incubated for 20 minutes in 10% normal donkey serum in PBS containing 0.1% triton x-100 and incubated overnight with 1:2500 rabbit anti-Gsx2 and 1:5000 guinea pig anti-Ascl1 (for the Gsx2:Ascl1 PLA) or 1:500 goat anti-Tcf3 and 1:5000 guinea pig anti-Ascl1 (for the Ascl1:Tcf3 PLA) at room temperature in a humidified chamber. The slides were then washed twice in PBS containing 0.1% triton X100, once with PBS without detergent and then once with 20mM Tris pH8, 150mM NaCl, 0.05% tween 20 (TBSt). Individual sections were then encircled using a hydrophobic pen before adding 100 µl of secondary antibody mix containing 20 µl of anti-rabbit +ve conjugated antibody from the PLA kit together with 5 µl −ve DNA conjugated anti-guinea pig antibody in dilution buffer containing 1% NDS and incubated at room temperature for two hours. Slides were washed twice in Olink buffer A (Millipore Sigma) and probe-ligation was initiated by addition of 0.5 µl ligase in 39.5 µl 1x ligase buffer and incubation at 37°C for 30 minutes. The slides were then washed 3 times in Olink buffer A and the amplification reaction was initiated by addition of 100 µl of polymerase to each section in 1x polymerase stock buffer and the slides were incubated overnight at 37°C in a humidified chamber. The following morning the slides were washed 3 times in TBS and briefly dipped in water. They were then coverslipped in Hoechst containing wet mounting medium and sealed with nail polish. PLA slides were imaged using an Olympus BX51 microscope with epifluorescence.

## Acknowledgements

We thank Ron Waclaw and Jeff Kuerbitz for critical reading of the manuscript. We also thank Jane Johnson (UT Southwestern, Dallas, TX) for providing the guinea pig anti-Ascl1 antibody and Hugo Bellen (Baylor College of Medicine, Houston Texas) for providing the *Ac-*Luciferase and Daughterless expression plasmids.

---

**Supplemental Figure 1.**
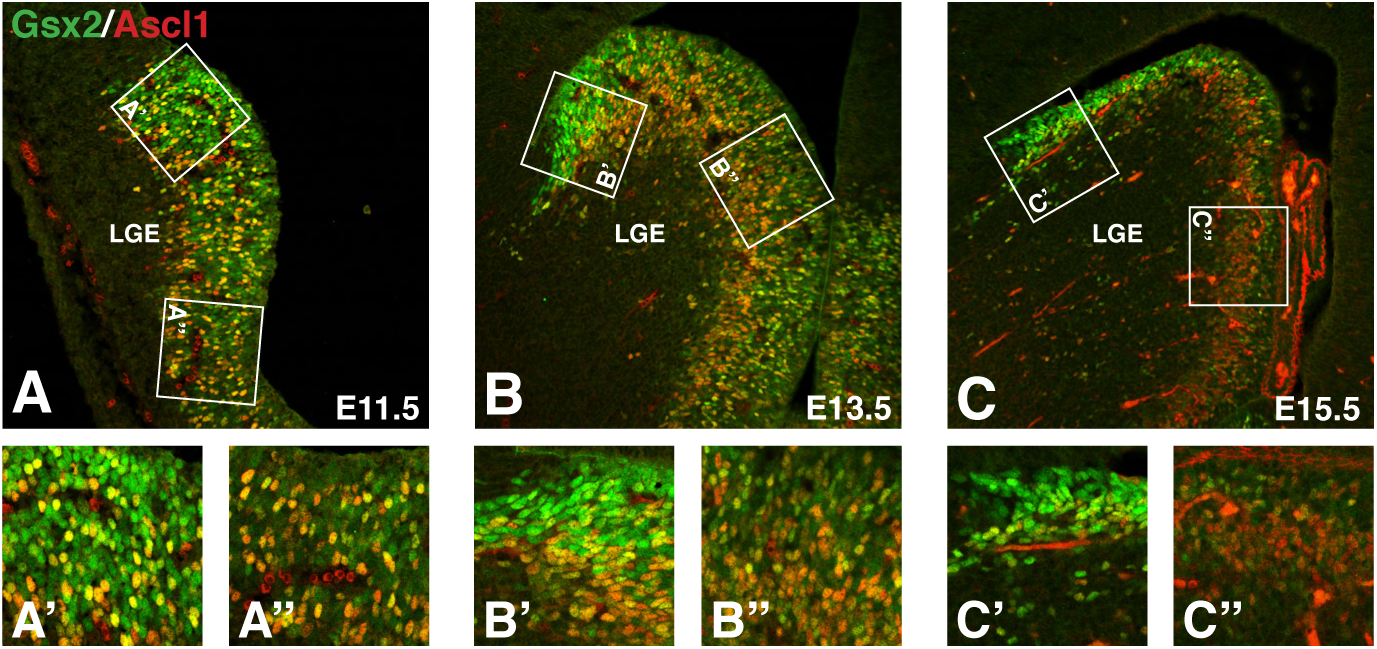
Co-expression of Gsx2 and Ascl1 in LGE progenitors over early neurogenic timepoints. Double immunohistochemistry for Gsx2 and Ascl1 in tissue sections from E11.5 (**A**), E13.5 (**B**) and E15.5 (**C**) mouse telencephalon. Boxed areas represent the VZ regions quantified for the dLGE (**A’, B’, C’**) and the vLGE (**A”, B”, C”**). Boxes have been rotated so that the apical surface is at the top of all images in A’, A”, B’, B”, C’ and C”. Results from the quantification are presented in Fig. 1**H, I**.

**Supplemental Figure 2.**
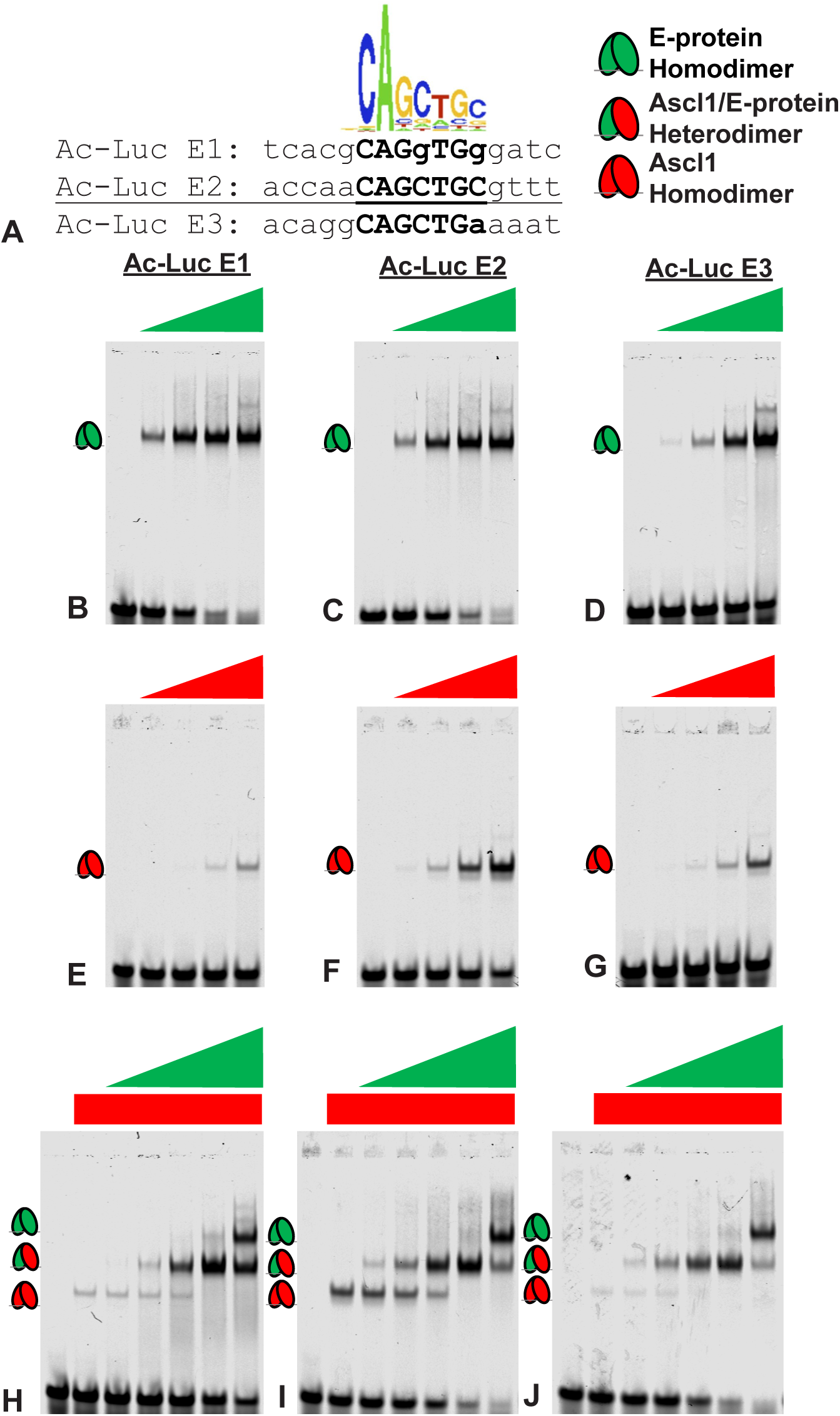
Ascl1 homodimers and heterodimers bind the 3 E-boxes from the Ac-luciferase reporter with varying affinities. **A**) Sequence logo for Ascl1 binding determined via ChIP on CHIP assay (Castro et al., 2011), and the E-box sequences from the *Ac-Luciferase* reporter used as probes in EMSAs. Matches to perfect Ascl1 binding site are capitalized. **B-D**) Binding of increasing levels of E-protein as E47-E47 homodimers to E-boxes from Ac-Luc **B**) E1, **C**) E2, and **D**) E3. **E-G**) Binding of increasing levels of Ascl1 as Ascl1-Ascl1 homodimers to E-boxes from Ac-Luc **E**) E1, **F**) E2, and **G**) E3. **H-J**) Binding of Ascl1 added at constant levels with increasing levels of E-protein as Ascl1-Ascl1 homodimers, Ascl1-E47 heterodimers, and E47-E47 homodimers to E-boxes from Ac-Luc **H**) E1, **I**) E2, and **J**) E3.

**Supplemental Figure 3.**
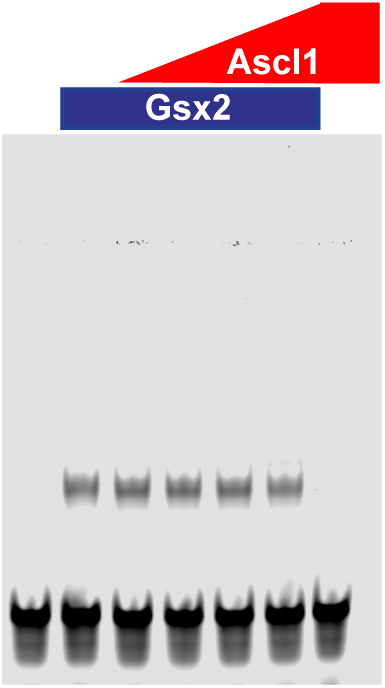
Ascl1 has no effect on Gsx2 DNA binding. Protein binding microarray data for mouse Gsx2 was used to generate a predicted, high affinity Gsx2 binding site (see Berger et al., 2008). EMSA shows binding of a constant amount of Gsx2 to a probe containing this high affinity site with increasing levels of Ascl1 added in each lane.

**Supplemental Table 1:** Plasmids generated for the experiements.

| **Insert** | **Amino Acids** | **Source** | **Destination**<br>**Vector** | **Integration**<br>**site** | **Method** |
| --- | --- | --- | --- | --- | --- |
| Gsx2 | 1-305 (Full length) | pBS SK- mouse clone | pGBKT7 | EcoR1-BamH1 | PCR |
| Gsx1 | 1-261 (Full length) | pBS SK- mouse clone | pGBKT7 | EcoR1-BamH1 | PCR |
| Ascl1 | 1-231 (Full length) | Embryonic mouse telencephalon RNA | pGADT7 | EcoR1-BamH1 | RT-PCR |
| Olig2 | 1-323 (Full length) | pCAGGS | pGADT7 | EcoR1-BamH1 | PCR |
| Ascl1 NT | 1-131 | pGADT7 FL clone | pGADT7 | EcoR1-BamH1 | PCR |
| Ascl1 bHLH | 97-193 | pGADT7 FL clone | pGADT7 | EcoR1-BamH1 | PCR |
| Ascl1 CT | 193-231 | pGADT7 FL clone | pGADT7 | EcoR1-BamH1 | PCR |
| Ascl1 Δ111-116 | $\Delta$ 111-116 | pGADT7 FL clone | pGADT7 | EcoR1-BamH1 | PCR |
| Ascl1 $\Delta$ 111-145 | $\Delta$ 111-145 | pGADT7 FL clone | pGADT7 | EcoR1-BamH1 | PCR |
| Ascl1 $\Delta$<br>145-150 | $\Delta$ 145-150 | pGADT7 FL clone | pGADT7 | EcoR1-BamH1 | PCR |
| Ascl1 $\Delta$<br>145-162 | $\Delta$ 145-162 | pGADT7 FL clone | pGADT7 | EcoR1-BamH1 | PCR |
| Ascl1 $\Delta$<br>145-191 | $\Delta$ 145-191 | pGADT7 FL clone | pGADT7 | EcoR1-BamH1 | PCR |
| Ascl1 $\Delta$<br>111-191 | $\Delta$ 111-191 | pGADT7 FL clone | pGADT7 | EcoR1-BamH1 | PCR |
| Ascl1 $\Delta$<br>111-179 | $\Delta$ 111-179 | pGADT7 FL clone | pGADT7 | EcoR1-BamH1 | PCR |
| Gsx2 Y3H<br>MCS1 | 1-305 (Full<br>length) | pCDNA6 FL clone | pBridge | EcoR1-BamH1 | PCR |
| TCF3 Y3H<br>MCS2 | 1-653 (isoform-1) | Mouse brain RNA | pBridge | Not1-BglII | RT-PCR |
| Gsx2 Y3H<br>MCS2 | 1-305 (Full<br>length) | pCDNA6 FL clone | pBridge | Not1-BglII | PCR |
| Tcf3 Y3H<br>MCS1 | 1-653 (isoform-1) | Mouse brain RNA | pBridge | EcoR1-BamH1 | RT-PCR |
| Tcf3 Y3H<br>MCS1,<br>Gsx2-MCS2 | 1-653 (isoform-1) | Mouse brain RNA | pBridge with<br>Gsx2 Y3H<br>MCS2 | EcoR1-BamH1 | PCR |
| Tcf3 Y3H<br>MCS1 | 1-653 (isoform-1) | Mouse brain RNA | pBridge with<br>Gsx2 Y3H<br>MCS2 | EcoR1-BamH1 | PCR |
| Ascl1 | 1-231 (Full<br>length) | pGADT7 clone | pAC5.1 | EcoRI-XhoI | Subcloning |
| Gsx2 | 1-305 (Full<br>length) | pCDNA6 FL clone | pAC5.1 | XhoI-XbaI | Subcloning |
| Gsx2 | N253A<br>1-305 (Full<br>length) | pGBKT7 clone | pAC5.1 | EcoRV-XhoI | Subcloning |
| Gsx2 | 167-305 | pGBKT7 clone | pET14b<br>(modified) | BamH1-NotI | PCR |
| Gsx2 | N253A<br>167-305 | pGBKT7 N253A<br>clone | pET14b<br>(modified) | BamH1-NotI | PCR |
| Ascl1 | 115-231 | pGADT7 clone | pET14b<br>(modified) | NdeI-KpnI | PCR |

**Supplemental Table 2:**
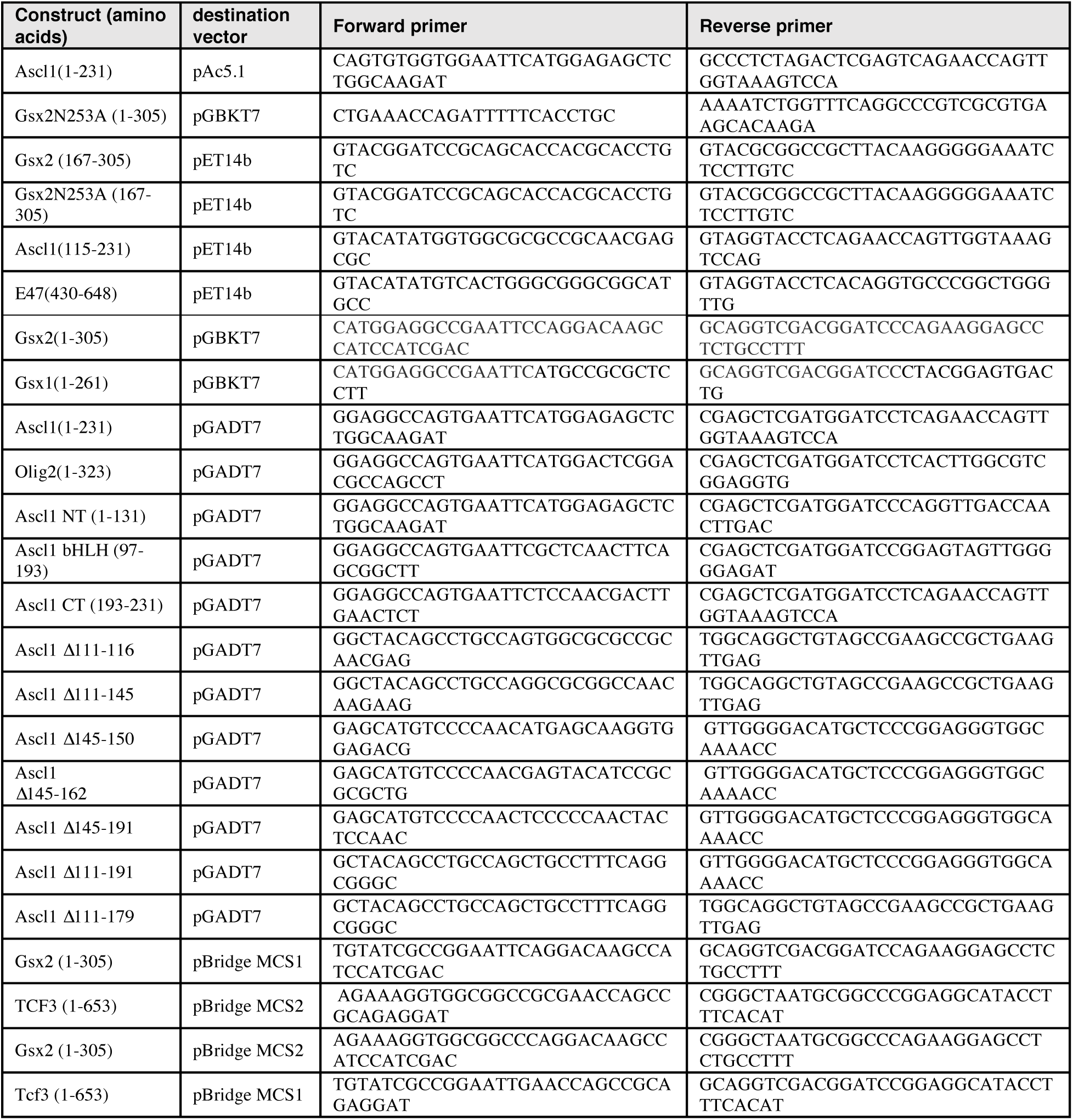
PCR primers used in cloning.

**Supplemental Table 3:** Contents of EMSAs in Figure 5.

| **Lane** | **Ascl1**<br>**(pmoles)** | **Tcf3**<br>**(pmoles)** | **Gsx2 N253A**<br>**(pmoles)** | **probe**<br>**(pmoles)** |
| --- | --- | --- | --- | --- |
| **1** | 0 | 0 | 0 | 0.04 |
| **2** | 2.5 | 0 | 0 | 0.04 |
| **3** | 2.5 | 0 | 20 | 0.04 |
| **4** | 2.5 | 0 | 40 | 0.04 |
| **5** | 2.5 | 0 | 80 | 0.04 |
| **6** | 0 | 0 | 0 | 0.04 |
| **7** | 2.5 | 0 | 0 | 0.04 |
| **8** | 0 | 0.078125 | 0 | 0.04 |
| **9** | 2.5 | 0.078125 | 0 | 0.04 |
| **10** | 2.5 | 0.078125 | 10 | 0.04 |
| **11** | 2.5 | 0.078125 | 20 | 0.04 |
| **12** | 2.5 | 0.078125 | 40 | 0.04 |
| **13** | 2.5 | 0.078125 | 80 | 0.04 |
| **14** | 0 | 0 | 0 | 0.04 |
| **15** | 0 | 0.3125 | 0 | 0.04 |
| **16** | 0.15625 | 0.3125 | 0 | 0.04 |
| **17** | 0.15625 | 0.3125 | 10 | 0.04 |
| **18** | 0.15625 | 0.3125 | 20 | 0.04 |
| **19** | 0.15625 | 0.3125 | 40 | 0.04 |
| **20** | 0.15625 | 0.3125 | 80 | 0.04 |
| **21** | 0 | 0 | 80 | 0.04 |
| **22** | 0 | 0 | 0 | 0.04 |
| **23** | 0.026 | 0 | 0 | 0.04 |
| **24** | 0 | 0.3125 | 0 | 0.04 |
| **25** | 0.026 | 0.3125 | 0 | 0.04 |
| **26** | 0.026 | 0.3125 | 20 | 0.04 |
| **27** | 0.026 | 0.3125 | 40 | 0.04 |
| **28** | 0.026 | 0.3125 | 80 | 0.04 |
| **29** | 0 | 0 | 0 | 0.04 |
| **30** | 0 | 0.3125 | 0 | 0.04 |
| **31** | 0 | 0.3125 | 20 | 0.04 |
| **32** | 0 | 0.3125 | 40 | 0.04 |
| **33** | 0 | 0.3125 | 80 | 0.04 |

## References

Bagchi, S., Fredriksson, R. and Wallén-Mackenzie Å. (2015). In Situ Proximity Ligation Assay (PLA). Methods. Mol. Biol.1318, 149–159.

Berger, M. F., Badis, G., Gehrke, A. R., Talukder, S., Philippakis, A. A., Peña-Castillo, L., Alleyne, T. M., Mnaimenh, S., Botvinnik, O. B., Chan, E. T. et al. (2008). Variation in homeodomain DNA binding revealed by high-resolution analysis of sequence preferences. Cell. 133, 1266–1276.

Bhide P. G. (1996). Cell cycle kinetics in the embryonic mouse corpus striatum. J. Comp. Neurol. 374, 506–522.

Castro, D. S., Martynoga, B., Parras, C., Ramesh, V., Pacary, E., Johnston, C., Drechsel, D., Lebel-Potter, M., Garcia, L. G., Hunt, C. et al. (2011). A novel function of the proneural factor Ascl1 in progenitor proliferation identified by genome-wide characterization of its targets. Genes. Dev. 25, 930–945.

Chapman, H., Waclaw, R. R., Pei, Z., Nakafuku, M. and Campbell, K. (2013). The homeobox gene Gsx2 controls the timing of oligodendroglial fate specification in mouse lateral ganglionic eminence progenitors. Development. 140, 2289–2298.

Chapman, H., Riesenberg, A., Ehrman, L. A., Kohli, V., Nardini, D., Nakafuku, M., Campbell, K. and Waclaw, R. R. (2018). Gsx transcription factors control neuronal versus glial specification in ventricular zone progenitors of the mouse lateral ganglionic eminence. Dev. Biol. 442, 115–126.

Chien, C. T., Bartel, P. L., Sternglanz, R. and Fields S. (1991). The two-hybrid system: a method to identify and clone genes for proteins that interact with a protein of interest. Proc. Natl. Acad. Sci U.S.A. 88, 9578–9582.

Corbin, J. G., Gaiano, N., Machold, R. P., Langston, A. and Fishell, G. (2000). The Gsh2 homeodomain gene controls multiple aspects of telencephalic development. Development. 127, 5007–5020.

Corbin, J. G., Rutlin, M., Gaiano, N. and Fishell, G. (2003). Combinatorial function of the homeodomain proteins Nkx2.1 and Gsh2 in ventral telencephalic patterning. Development. 130, 4895–906.

Deacon, T.W., Pakzaban, P. and Isacson, O. (1994). The lateral ganglionic eminence is the origin of cells committed to striatal phenotypes: neural transplantation and developmental evidence. Brain. Res. 668, 211–219.

Fietz, S. A., Kelava, I., Vogt, J., Wilsch-Bräuninger, M., Stenzel, D., Fish, J. L., Corbeil, D., Riehn, A., Distler, W., Nitsch, R. et al. (2010). OSVZ progenitors of human and ferret neocortex are epithelial-like and expand by integrin signaling. Nat. Neurosci. 13, 690–699.

Fish, J. L., Dehay, C., Kennedy, H. and Huttner, W. B. (2008). Making bigger brains-the evolution of neural-progenitor-cell division. J. Cell. Sci. 121, 2783–2793.

Fode, C., Ma, Q., Casarosa, S., Ang, S. L., Anderson, D. J. and Guillemot, F. (2000). A role for neural determination genes in specifying the dorsoventral identity of telencephalic neurons. Genes. Dev. 14, 67–80.

Fogarty, M., Grist, M., Gelman, D., Marín, O., Pachnis, V. and Kessaris, N. (2007). Spatial genetic patterning of the embryonic neuroepithelium generates GABAergic interneuron diversity in the adult cortex. J. Neurosci. 27, 10935–10946.

Gerfen, C. R. (1992). The neostriatal mosaic: multiple levels of compartmentalization in the basal ganglia, Annu. Rev. Neurosci. 15, 285–320.

Graybiel, A. M. and Ragsdale, C. W. Jr. (1978). Histochemically distinct compartments in the striatum of human, monkeys, and cat demonstrated by acetylcholinesterase staining. Proc. Natl. Acad. Sci. USA 75, 5723–5726.

Hanashima, C., Shen, L., Li, S. C. and Lai, E. (2002). Brain factor-1 controls the proliferation and differentiation of neocortical progenitor cells through independent mechanisms. J. Neurosci. 22, 6526–6536.

Hansen, D. V., Lui, J. H., Parker, P. R. and Kriegstein, A. R. (2010). Neurogenic radial glia in the outer subventricular zone of human neocortex. Nature. 464, 554–561.

Haubensak, W., Attardo, A., Denk, W. and Huttner, W. B. (2004). Neurons arise in the basal neuroepithelium of the early mammalian telencephalon: a major site of neurogenesis. Proc. Natl. Acad. Sci U.S.A. 101, 3196–3201.

Henke, R. M., Meredith, D. M., Borromeo, M. D., Savage, T. K. and Johnson, J. E. (2009). Ascl1 and Neurog2 form novel complexes and regulate Delta-like3 (Dll3) expression in the neural tube. Dev. Biol. 328, 529–540.

Hinds J. W. (1968) Autoradiographic study of histogenesis in the mouse olfactory bulb. I. Time of origin of neurons and neuroglia. J. Comp. Neurol. 134, 287–304.

Imayoshi, I., Isomura, A., Harima, Y., Kawaguchi, K., Kori, H., Miyachi, H., Fujiwara, T., Ishidate, F. and Kageyama, R. (2013). Oscillatory control of factors determining multipotency and fate in mouse neural progenitors. Science. 342, 1203–1208.

Jafar-Nejad, H., Acar, M., Nolo, R., Lacin, H., Pan, H., Parkhurst, S. M. and Bellen, H. J. (2003). Senseless acts as a binary switch during sensory organ precursor selection. Genes. Dev. 17, 2966–2978.

Johnson J. E., Birren, S. J., Saito, T. and Anderson, D. J. (1992). DNA binding and transcriptional regulatory activity of mammalian achaete-scute homologous (MASH) proteins revealed by interaction with a muscle-specific enhancer. Proc. Natl. Acad. Sci. 89, 3596–3600.

Johnston, J. G., Gerfen, C. R., Haber, S. N. and van der Kooy, D. (1990). Mechanisms of striatal pattern formation: conservation of mammalian compartmentalization. Brain. Res. Dev. Brain. Res. 57, 93–103.

Kessaris N., /Fogarty, M., Iannarelli, P., Grist, M., Wegner, M. and Richardson, W. D. (2006). Competing waves of oligodendrocytes in the forebrain and postnatal elimination of an embryonic lineage. Nat. Neurosci. 9, 173–179.

Kelly, S. M., Raudales, R., He, M., Lee, J. H., Kim, Y., Gibb, L. G., Wu, P., Matho, K., Osten, P., Graybiel, A. M. et al. (2018). Radial Glial Lineage Progression and Differential Intermediate Progenitor Amplification Underlie Striatal Compartments and Circuit Organization. Neuron. 99, 345–361.

Kuerbitz, J., Arnett, M., Ehrman, S., Williams, M. T., Vorhees, C. V., Fosher, S. E., Garratt, A. N., Muglia, L. J., Waclaw, R. R. and Campbell, K. (2018). Loss of Intercalated Cells (ITCs) in the Mouse Amygdala of Tshz1 Mutants Correlates with Fear, Depression, and Social Interaction Phenotypes. J. Neurosci. 38, 1160–1177.

Letteboer, S. J. and Roepman, R. (2008). Versatile screening for binary protein-protein interactions by yeast two-hybrid mating. Methods. Mol. Biol. 484, 145–159.

Lin J. S. and Lai E. M. (2017). Protein-Protein Interactions: Co-Immunoprecipitation. Methods. Mol. Biol. 1615, 211–219.

Lindtner, S., Catta-Preta, R., Tian, H., Su-Feher, L., Price, D. J., Dickel, D. E., Greiner, V., Silberberg, S. N., McKinsey, G. L., McManus, M. T. et al. (2019). Genomic Resolution of DLX-Orchestrated Transcriptional Circuits Driving Development of Forebrain GABAergic Neurons. Cell. Rep. 28, 2048–2063.

Long, J. E., Cobos, I., Potter, G. B. and Rubenstein, J. L. (2009a). Dlx1&2 and Mash1 transcription factors control MGE and CGE patterning and differentiation through parallel and overlapping pathways. Cereb. Cortex. Suppl 1, i96–106.

Long, J. E., Swan, C., Liang, W. S., Cobos, I., Potter, G. B. and Rubenstein, J. L. (2009b). Dlx1&2 and Mash1 transcription factors control striatal patterning and differentiation through parallel and overlapping pathways. J. Comp. Neurol. 512, 556–572.

López-Juárez, A., Howard, J., Ullom, K., Howard, L., Grande, A., Pardo, A., Waclaw, R., Sun, Y. Y., Yang, D., Kuan, C. Y. et al. (2013). Gsx2 controls region-specific activation of neural stem cells and injury-induced neurogenesis in the adult subventricular zone, Genes. Dev. 27, 1272–1287.

Massari M. E. and Murre C. (2000). Helix-loop-helix proteins: regulators of transcription in eucaryotic organisms. Mol. Cell. Biol. 20, 429–440.

Méndez-Gómez H. R. and Vicario-Abejón C. (2012). The homeobox gene Gsx2 regulates the self-renewal and differentiation of neural stem cells and the cell fate of postnatal progenitors. PLoS. One. 7, e29799.

Nakada, Y., Hunsaker, T. L., Henke, R. M. and Johnson, J. E. (2004). Distinct domains within Mash1 and Math1 are required for function in neuronal differentiation versus neuronal cell-type specification. Development. 131, 1319–1330.

Noctor, S. C., Martínez-Cerdeño, V., Ivic, L. and Kriegstein, A. R. (2004). Cortical neurons arise in symmetric and asymmetric division zones and migrate through specific phases. Nat. Neurosci. 7, 136–144.

Olsson, M., Campbell, K., Wictorin, K. and Björklund, A. (1995). Projection neurons in fetal striatal transplants are predominantly derived from the lateral ganglionic eminence. Neuroscience. 69, 1169–1182.

Olsson, M., Björklund, A. and Campbell, K. (1998). Early specification of striatal projection neurons and interneuronal subtypes in the lateral and medial ganglionic eminence. Neuroscience. 84, 867–876.

Pei, Z., Wang, B., Chen, G., Nagao, M., Nakafuku, M. and Campbell, K. (2011). Homeobox genes Gsx1 and Gsx2 differentially regulate telencephalic progenitor maturation. Proc. Natl. Acad. Sci U.S.A. 108, 1675–1680.

Pilz, G. A., Shitamukai, A., Reillo, I., Pacary, E., Schwausch, J., Stahl, R., Ninkovic, J., Snippert, H. J., Clevers, H., Godinho, L. et al. (2013). Amplification of progenitors in the mammalian telencephalon includes a new radial glial cell type. Nat. Commun. 4, 2125

Pla, R., Stanco, A., Howard, M. A., Rubin, A. N., Vogt, D., Mortimer, N., Cobos, I., Potter, G. B., Lindtner, S., Price, J. D., et al. (2018). Dlx1 and Dlx2 Promote Interneuron GABA Synthesis, Synaptogenesis, and Dendritogenesis. Cereb. Cortex. 28, 3797–3815.

Qin, S., Madhavan, M., Waclaw, R. R., Nakafuku, M. and Campbell, K. (2016). Characterization of a new Gsx2-cre line in the developing mouse telencephalon. Genesis. 54, 542–549.

Qin, S., Ware, S. M., Waclaw, R. R. and Campbell, K. (2017). Septal contributions to olfactory bulb interneuron diversity in the embryonic mouse telencephalon: role of the homeobox gene Gsx2. Neural. Dev. 12:13.

Smart, I. H. (1976). A pilot study of cell production by the ganglionic eminences of the developing mouse brain. J. Anat. 121, 71–84.

Söderberg, O., Gullberg, M., Jarvius, M., Ridderstråle, K., Leuchowius, K. J., Jarvius, J., Wester, K., Hydbring, P., Bahram, F., Larsson, L. G. et al. (2006). Direct observation of individual endogenous protein complexes in situ by proximity ligation. Nat. Methods. 3, 995–1000.

Stenman, J., Toresson, H. and Campbell, K. (2003). Identification of two distinct progenitor populations in the lateral ganglionic eminence: implications for striatal and olfactory bulb neurogenesis. J. Neurosci. 23, 167–174.

Szucsik, J. C., Witte, D. P., Li, H., Pixley, S. K., Small, K. M. and Potter, S. S. (1997). Altered forebrain and hindbrain development in mice mutant for the Gsh-2 homeobox gene. Dev. Biol. 191, 230–242.

Takebayashi, H., Yoshida, S., Sugimori, M., Kosako, H., Kominami, R., Nakafuku, M. and Nabeshima, Y. (2000). Dynamic expression of basic helix-loop-helix Olig family members: implication of Olig2 in neuron and oligodendrocyte differentiation and identification of a new member, Olig3. Mech. Dev. 99, 143–148.

Tirode, F., Malaguti, C., Romero, F., Attar, R., Camonis, J. and Egly, J. M. (1997). A conditionally expressed third partner stabilizes or prevents the formation of a transcriptional activator in a three-hybrid system. J. Biol. Chem. 272, 22995–22999.

Toresson, H., Potter, S. S. and Campbell, K. (2000). Genetic control of dorsal-ventral identity in the telencephalon: opposing roles for Pax6 and Gsh2. Development. 127, 4361–4371.

Toresson, H. and Campbell, K. (2001). A role for Gsh1 in the developing striatum and olfactory bulb of Gsh2 mutant mice. Development. 128, 4769–4780.

Tucker, E. S., Polleux, F. and LaMantia, A. S. (2006). Position and time specify the migration of a pioneering population of olfactory bulb interneurons. Dev. Biol. 297, 387–401.

Ueki, Y., Wilken, M. S., Cox, K. E., Chipman, L., Jorstad, N., Sternhagen, K., Simic, M., Ullom, K., Nakafuku, M. and Reh, T. A. (2015). Transgenic expression of the proneural transcription factor Ascl1 in Müller glia stimulates retinal regeneration in young mice. Proc. Natl. Acad. Sci U.S.A. 112, 13717–13722.

Uhl, J. D., Cook, T. A. and Gebelein, B. (2010). Comparing anterior and posterior Hox complex formation reveals guidelines for predicting cis-regulatory elements. Dev. Biol. 343, 154–166.

Uhl, J. D., Zandvakili, A. and Gebelein, B. (2016). A Hox transcription factor collective binds a highly conserved Distal-less cis-regulatory module to generate robust transcriptional outcomes. PLoS. Genet. 12, e1005981.

van der Kooy D. and Fishell G. (1987). Neuronal birthdate underlies the development of striatal compartments. Brain. Res. 401, 155–161.

Waclaw, R. R., Allen, Z. J., 2nd, Bell, S. M., Erdélyi, F., Szabó. G., Potter, S. S. and Campbell, K. (2006). The zinc finger transcription factor Sp8 regulates the generation and diversity of olfactory bulb interneurons. Neuron. 49, 503–516.

Waclaw, R. R., Wang, B., Pei, Z., Ehrman, L. A. and Campbell, K. (2009). Distinct temporal requirements for the homeobox gene Gsx2 in specifying striatal and olfactory bulb neuronal fates. Neuron. 63, 451–465.

Waclaw, R. R., Ehrman, L. A., Pierani, A. and Campbell, K. (2010). Developmental origin of the neuronal subtypes that comprise the amygdalar fear circuit in the mouse. J. Neurosci. 30, 6944–6953.

Wang, B., Waclaw, R. R., Allen, Z. J., 2nd, Guillemot, F. and Campbell, K. (2009). Ascl1 is a required downstream effector of Gsx gene function in the embryonic mouse telencephalon. Neural. Dev. 4:5.

Wang, B., Long, L. E., Flandin, P., Pla, R., Waclaw, R. R., Campbell, L. and Rubenstein, J. L. (2013). Loss of Gsx1 and Gsx2 function rescues distinct phenotypes in Dlx1/2 mutants. J. Comp. Neurol. 521, 1561–1584.

Wichterle, H., Turnbull, D. H., Nery, S., Fishell, G. and Alvarez-Buylla, A. (2001). In utero fate mapping reveals distinct migratory pathways and fates of neurons born in the mammalian basal forebrain. Development. 128, 3759–3771.

Wilsch-Bräuninger, M., Florio, M. and Huttner, W. B. (2016). Neocortex expansion in development and evolution - from cell biology to single genes. Curr. Opin. Neurobiol. 39, 122–132.

Winterbottom, E. F., Illes, J. C., Faas, L. and Isaacs H. V. (2010). Conserved and novel roles for the Gsh2 transcription factor in primary neurogenesis. Development. 137, 2623–31.

Winterbottom, E. F., Ramsbottom, S. A., Isaacs, H. V. (2011). Gsx transcription factors repress Iroquois gene expression. Dev. Dyn. 240, 1422–1429.

Yun, K., Potter, S. and Rubenstein, J. L. (2001). Gsh2 and Pax6 play complementary roles in dorsoventral patterning of the mammalian telencephalon. Development. 128, 193–205.

Yun, K., Fischman, S., Johnson, J., Hrabe de Angelis, M., Weinmaster, G. and Rubenstein, J. L. (2002). Modulation of the notch signaling by Mash1 and Dlx1/2 regulates sequential specification and differentiation of progenitor cell types in the subcortical telencephalon. Development. 129, 5029–5040.

Yun, K., Garel, S., Fischman, S. and Rubenstein, J. L. (2003). Patterning of the lateral ganglionic eminence by the Gsh1 and Gsh2 homeobox genes regulates striatal and olfactory bulb histogenesis and the growth of axons through the basal ganglia. J. Comp. Neurol. 461, 151–165.

Zhang, X., McGrath, P. S., Salomone, J., Rahal, M., McCauley, H. A., Schweitzer, J., Kovall, R., Gebelein, B. and Wells, J. M. (2019). A Comprehensive Structure-Function Study of Neurogenin3 Disease-Causing Alleles during Human Pancreas and Intestinal Organoid Development. Dev. Cell. 50, 367–380.

